# Altered relaxation and Mitochondria-Endoplasmic Reticulum Contacts Precede Major (Mal)adaptations in Aging Skeletal Muscle and are Prevented by Exercise

**DOI:** 10.1101/2025.01.14.633043

**Authors:** Ryan J. Allen, Ana Kronemberger, Qian Shi, Marshall Pope, Elizabeth Cuadra-Muñoz, Wangkuk Son, Long-Sheng Song, Ethan J. Anderson, Renata O. Pereira, Vitor A. Lira

## Abstract

Sarcopenia, or age-related muscle dysfunction, contributes to morbidity and mortality. Besides decreases in muscle force, sarcopenia is associated with atrophy and fast-to-slow fiber type switching, which is typically secondary to denervation in humans and rodents. However, very little is known about cellular changes preceding these important (mal)adaptations. To this matter, mitochondria and the sarcoplasmic reticulum are critical for tension generation in myofibers. They physically interact at the boundaries of sarcomeres forming subcellular hubs called mitochondria-endo/sarcoplasmic reticulum contacts (MERCs). Yet, whether changes at MERCs ultrastructure and proteome occur early in aging is unknown. Here, studying young adult and older mice we reveal that aging slows muscle relaxation leading to longer excitation-contraction-relaxation (ECR) cycles before maximal force decreases and fast-to-slow fiber switching takes place. We reveal that muscle MERC ultrastructure and mitochondria-associated ER membrane (MAM) protein composition are also affected early in aging and are closely associated with rate of muscle relaxation. Additionally, we demonstrate that regular exercise preserves muscle relaxation rate and MERC ultrastructure in early aging. Finally, we profile a set of muscle MAM proteins involved in energy metabolism, protein quality control, Ca^2+^ homeostasis, cytoskeleton integrity and redox balance that are inversely regulated early in aging and by exercise. These may represent new targets to preserve muscle function in aging individuals.

## Introduction

By 2050, individuals aged 60 years or older will represent over 20 % of the global population (Chatterji et al, 2015). As this demographic expands, so do the inevitable ailments accompanying biological age. One such condition is sarcopenia, defined as the progressive and generalized loss of muscle mass and function with age, which reportedly affects up to 29 % of older individuals (Cruz-Jentoft et al, 2014; Lang et al, 2010). Sarcopenia not only detracts significantly from quality of life but also markedly increases risks of morbidity and mortality (Budui et al, 2015). Therefore, healthcare efforts to attenuate the progression of sarcopenia (herein referred to as age-related muscle dysfunction) are critical for improving the health span of our aging population. However, effective therapies remain limited because our understanding of the structural and functional events occurring early in muscle aging remains scarce.

It is well documented that muscle maximal force decreases dramatically in advanced ages (Doherty, 2003; Lang et al, 2010; Sayer et al, 2024). At the cellular level, these functional changes are accompanied by pronounced atrophy of type 2 fibers and involve fast-to-slow (i.e., type 2-to-type 1) fiber type switching, which is typically secondary to type 2 fiber denervation (Aagaard et al, 2010; Dowling et al, 2023; Lang et al, 2010; Larsson et al, 2019). However, very little is known about functional changes preceding these important (mal)adaptations. Though it is well recognized that regular exercise alleviates the impact of aging on skeletal muscle, exercise-mediated changes that occur early in aging is not well-characterized (Budui et al, 2015; Cartee et al, 2016; Schumann et al, 2022; Wang et al, 2022).

Both the sarcoplasmic reticulum (SR), a specialized form of the endoplasmic reticulum (ER) in muscle, and mitochondria are required for tension generation in skeletal myofibers. Mitochondria-endo/sarcoplasmic reticulum contacts (MERCs), also known as mitochondria-associated ER/SR membranes (MAMs), are sites of physical coupling between the outer mitochondrial membrane (OMM) and the ER/SR. Across cell populations, MERCs modulate numerous processes such as ion and nucleotide exchange, lipid metabolism, and autophagosome formation, among others (Giacomello et al, 2016; Hamasaki et al, 2013). Within muscle, MERCs are predominately located at sarcomeres near the intersection of the I-band and the Z-disc. Because mitochondria and the ER/SR become dysfunctional in sarcopenic muscle and adapt to exercise, MERCs likely play a role in mediating these changes (Baehr et al, 2016; Bohnert et al, 2018; Kauppila et al, 2017; Kubat et al, 2023; Paez et al, 2023; Parry et al, 2020; Tarnopolsky et al, 2007) In fact, recent studies have documented irregular MERC/MAM morphology and protein composition in skeletal muscle across multiple conditions (Hinton et al, 2022; Lu et al, 2022; Thoudam et al, 2018; Tubbs et al, 2018).

In this study, we reveal that early muscle aging is characterized by slowed relaxation and increased fatigue. This occurs before reductions in muscle force or changes to fiber type and is effectively reversed by regular exercise. Additionally, we demonstrate that muscle MERC/MAM ultrastructure and protein composition are affected at this stage and are closely associated with the changes to muscle relaxation. Moreover, we reveal for the first time a set of muscle MAM proteins that are inversely regulated by aging and exercise offering novel insights into potential therapeutic targets to preserve or rescue muscle function in our aging population.

## Experimental Procedures

### Animals

All experimental procedures were conducted using male, C57BL6N mice and were approved by the University of Iowa Institutional Animal Care and Use Committee. Animals were housed onsite (Medical Laboratories Vivarium, University of Iowa, Iowa City, IA) in 21°C controlled rooms operating on 12-hour light-dark cycles. Mice had ad-lib access to water and Teklad global soy protein-free, irradiated diet (2920x). Our experimental groups consisted of young (HYA, 6 months), old (eAMD, 21 months), and very old (aAMD, 31 months) animals. The exercise group (eAMD+Ex) started training at 19 months of age, finishing the intervention at the same age as eAMD. Euthanasia was performed via cervical dislocation following anesthetization by 3 % isofluorane gas. Hindlimb muscles were harvested at the beginning of the light cycle under basal conditions and at least 36 hours after the last bout of exercise.

### *In vivo* Muscle function

Set-up, assessment, and analysis of muscle force using the 1300A 3-in-1 Whole Animal System (1300A; Aurora Scientific, Aurora, ON, Canada) was performed as previously described (Fuqua et al, 2019). Each animal was first subjected to a maximal stimulus (150Hz), and then the fatigue protocol was initiated after a 90-second rest period. The fatigability protocol includes 70 submaximal tetanic contractions (50 Hz) with 5 sec between each. Percentage of force preservation was calculated by using the mean of the last 5 contractions as a percentage of the first 5 contractions during the fatigue test. The DMA software automatically reports other variables (i.e., max rate of contraction/relaxation), which were also included in this study. Mean rate of relaxation represents the average rate observed during the fatigability protocol. Plantarflexor (gastrocnemius (GA), soleus (SOL), and plantaris (PL)) and dorsiflexor (tibialis anterior (TA) and extensor digitorum longus (EDL)) muscles were harvested no earlier than 48 hours following the last functional test.

### Treadmill intervention protocol and exercise capacity testing

The 6-8 week intervention was performed on a treadmill (Panlab/ Harvard Apparatus 5- lane Touchscreen Treadmill) using a separate group of age-matched eAMD mice (i.e., regular exercised from 19 to 21 months of age). Training began at 5 % incline and speed of 12 m/min (i.e., ∼50 % VO_2_max (Schefer et al, 1996)) for 25 minutes. Animals were trained 5 days per week at the beginning of the light cycle as recently described (Schenk et al, 2024). In short, running time was increased by one minute at each session up to a maximum of 50 minutes, and the speed was increased by 2 m/min every week. All sessions after the first week were at 10 % incline. Fitness levels were maintained by using the volume of the final training day until the completion of post-intervention testing. Tissues were harvested no earlier than 48 hours following the last bout of exercise.

Treadmill exhaustion tests were performed to quantify exercise capacity. Baseline and control animals were given a 3-day acclimation period of 6 m/min for 10 min. Following a 5- minute warm-up at 6 m/min, the exhaustion protocol began at 7.2 m/min with a 5 % incline and acceleration of 0.27 m/min until failure. Mice were considered exhausted after falling three times to the electric grid at the rear portion of the treadmill within 10 seconds. Total work capacity was calculated and reported based on running distance and body weight.

### Mitochondrial O_2_ consumption, ATP production, and H_2_O_2_ emission in permeabilized fibers

Preparation and handling of saponin-permeabilized mouse red GA fibers has been previously described in detail (Lark et al, 2016). Additional information on buffers and reagents can be found within the Supplementary Document 8. Both ATP production and high-resolution respirometry were conducted using the Oroboros Oxygraph-2k with a previous protocol from our group (O_2_k; Innsbruck, Austria) (Harris et al, 2022; Turner et al, 2022). Both ATP production and O_2_ consumption were normalized to wet tissue weight.

Hydrogen peroxide (H_2_O_2_) emission in permeabilized red GA fibers was measured with Amplex Red reagent as previously described (Anderson et al, 2007). Specific information can be found within the Supplementary Document 8. Rates of H_2_O_2_ emission before and after the addition of succinate were calculated via the slope of ΔF/min after subtracting background and was normalized to wet fiber weight.

### GSH/GSSG measurement in muscle

Colorimetric determination of total glutathione, reduced glutathione (GSH), oxidized glutathione (GSSG) and GSH/GSSG was assayed in a microplate reader (Biotek Epoch Microplate Spectrophotometer, Winooski, Vermont). For total glutathione, ∼5 mg of frozen GA powder was homogenized in 150 µL of TEE buffer (10 mM Tris, 1 mM EDTA, 1 mM EGTA, 0.5 % Tween-20, pH 7.4). For GSSG assessment, samples followed the same procedure but included more tissue (∼15 mg) and were homogenized in 0.3 mM Methyl-2-inylpyridinium (M_2_VP) in TEE buffer. Homogenates were centrifuged at 10,000 x *g* for 10 minutes, and pellets were discarded. Next, protein concentration was determined for each sample using the Pierce 660nm Protein Assay. To fit within the standard curve, GSH samples were diluted 10-fold. A consistent amount of protein/sample was added to each well in a volume of 25 µL. Standards of 0.05 µM, 0.125 µM, 0.25 µM, 0.75 µM, and 1.5 µM were run in triplicates with either TEE buffer (GSH) or TEE buffer + M_2_VP (GSSG). Glutathione Reductase (10 U/mL) mixed with DTNB (1 mM) (1:1 ratio), totaling 50 µL, was added to each well and incubated at room temperature for 10 min. Finally, 25 µL of NADPH was added to each well and immediately placed into the microplate reader. The reaction was read at 405 nm every minute for 6 min. Total glutathione and GSSG were reported based on the concentration of the samples in relation to their respective standard curves. GSH was determined by subtracting GSSG from the total glutathione concentration.

### Calcium uptake in isolated mitochondria

Excised TA muscles were weighed and placed on ice cold mitochondria isolation medium (MIM) (300mM Sucrose, 10mM Na-HEPES, 0.2mM EDTA, 1mM EGTA, pH 7.2 adjusted with glacial acetic acid). The tissue was finely minced and digested with 10mL trypsin (125ug/mL in MIM) on ice for two minutes. Following addition of 10mL trypsin inhibitor (650ug/mL in MIM + BSA [1mg/mL]), tissue was centrifuged at 600xg for 10 min at 4°C. The supernatant was collected and homogenized with Teflon pestles in Potter-Elvehjem glass tubes using a Wheaton Overhead Stirrer. Homogenates were then centrifuged at 800xg for 10 min at 4°C. The supernatant was collected and further centrifuged at 8000xg for 15 min at 4°C. This supernatant was discarded, and the mitochondrial pellet was cleaned via aspiration of contaminants. Another round of centrifugation was repeated, and the final mitochondrial pellet was resuspended in a small volume of MIM.

Calcium retention capacity (CRC) and maximal uptake speed in isolated mitochondria were measured using Calcium green-5N, as previously described (Murphy et al, 1996). Fluorescence was measured at 37°C in the BMG LABTECH (BMG Labtech, Cary, NC) microplate reader following 5µL injection of 0.6mM CaCl2 per well every 42 seconds. CRC was calculated based on the total amount of free Ca2+ ingestion until the fluorescent signal became saturated (i.e., opening of mitochondrial permeability transition pore [mPTP]). Maximal calcium uptake speed was quantified by the highest rate of free calcium uptake achieved during a single injection.

### TEM sample preparation, acquisition, and analysis

Harvested GA muscles were immediately weighed, placed into cold saline and separated into red and white components. Red GAs were fixed in 2.5 % glutaraldehyde and 1 % paraformaldehyde (Boudina et al, 2007). Post-fixation, embedding, sectioning, and imaging were conducted at the University of Iowa Microscopy Core Facility. Images at 2000 X and 8000 X magnifications for mitochondrial and MERC measurements, respectively, were obtained using a Hitachi HT7800 transmission electron microscope (Hitachi, Tokyo, Japan). Following the blinding of experimental groups, all morphological parameters were quantified using ImageJ as previously described (Lam et al, 2021). In short, mitochondria morphological measurements (i.e., mitochondrion area, circularity, and density) were assessed from 2000 X images using the free hand tool to encircle all of the individual intermyofibrillar mitochondria. The line tool was used to measure the MERC length (i.e., lengths of individual SR membranes that were within 50 nm of the OMM) and MERC width (i.e., the closest distance between the cytosolic faces of the SR and OMM). MERC coverage was quantified by determining the cumulative distance of MERC length(s) at an individual mitochondrion in relation to that same mitochondrion’s circumference and expressed as a percentage.

### Histological analysis

For all histological analyses, dissected plantares were quickly pinned and frozen in liquid nitrogen-cooled isopentane. Prior to sectioning, frozen muscles were embedded in optimal cutting temperature compound. Cross-sectional 5 µm serial sections were obtained using a Leica CM1520 cryostat (Leica Biosystems, Deer Park, IL). All slides were imaged on an Olympus BX63 microscope (Olympus, Tokyo, Japan). To assess fiber type and fiber diameter, sections were post-fixed in acetone and blocked with 5 % normal horse serum in 1X PBST. Next, slides were incubated in primary antibodies overnight at 4°C: anti-MyHC 1/2A/2B (BA-F8/SC71/BF-F3 from Developmental Studies Hybridoma Bank, Iowa City, IA) and anti-Laminin (L9393 from Sigma-Aldrich). The following day, slides were incubated for 30 minutes in combined secondary antibodies: 647 nm IgG2B (A21242), 555 nm IgG1 (A21127), 488 nm IgM (A21042), and 405 nm IgG (A31556B). Three washes in 1X PBST preceded each of the steps. Finally, coverslips were mounted on each slide using Vectashield Plus Antifade Mounting Medium (H-1900, Vector Labs, Newark, CA). Analysis of minimum Feret Diameter and fiber composition was performed on ImageJ software, as previously described (Turner et al, 2022). On average, 350 myofibers were inspected per section.

### Isolation of mitochondria-associated ER membranes (MAMs)

Subcellular MAM fractions were obtained from the GA muscle using a previously described protocol in muscle (Lu et al, 2022; Wieckowski et al, 2009). Additional details can be found with the Supplementary Document 8. For every fractionation, both GAs were used. Protein concentration was determined using the Pierce 660-nm Protein Assay.

### Immunoblot analysis

Whole GA lysates were prepared for SDS-PAGE as previously described (Fuqua et al, 2019). Isolated MAM fractions were prepared for SDS-PAGE by mixing the final samples (suspended in MRB) with 4X Loading Dye (50 mM Tris·HCl, pH 6.8, 1 % sodium dodecyl sulfate (SDS), 10 % glycerol, 20 mM dithiothreitol, 127 mM 2-mercaptoethanol, and 0.01 % bromophenol blue) in a 3:1 ratio and denatured at 95°C for 5 minutes. The primary antibodies used for this study were: VDAC1 (Cell Signaling, D73D12), Alpha-tubulin (Abcam, ab7291), PERK (C33E10), Citrate Synthase (Sigma, sab2701077) ASCL4 (Santa Cruz, sc-365230), OXPHOS antibody cocktail (Abcam, ab110413), and COX IV (Abcam, ab33985). PVDF membranes were used for all immunoblot analysis and images were taken in the Li-Cor Odyssey CLX system (LI-COR Biosciences, Lincoln, NE, USA). Protein signals were normalized to Ponceau staining.

### TMT sample preparation

Supplementary document 8 outlines sample clean-up and digestion. This was followed by LC/MS-MS to normalize samples based upon spectral counting (Erdjument-Bromage et al, 2018; Poston et al, 2013; Rappsilber et al, 2007). TMT labeling was conducted as previously described (Zecha et al, 2019). Essentially, equivalent aliquots were reconstituted in a 17.5 µL of triethylammonium bicarbonate (TEAB). Samples were mixed 2:1 w/w with TMT-10-plex labeling reagent from anhydrous acetonitrile. Reactions continued for 1 hour at room temperature and were then quenched with hydroxyl amine. Once barcoded, the samples were combined. The pooled sample was pre-fractionated into 8 fractions and a pass-through fraction which contained the hydrolyzed reagents and dimers using a Pierce High pH Reversed-Phase Peptide Fractionation Kit (Thermo Scientific,84868) according to the manufacturer’s instructions.

### Proteomic data acquisition and search

Data acquisitions on the LUMOS began with a survey scan (m/z 380 -1800) acquired in the Orbitrap at a resolution of 120,000 (MS1) with Automatic Gain Control (AGC) set to 3E06 and a maximum injection time of 50 ms. MS1 scans are acquired every 3 s during the 160 min gradient. The most abundant precursors were selected among 2-6 charge state ions at a 1E05 AGC and 70 ms maximum fill time. Ions were isolated with a 0.6 Th window using the multi-segment quadrupole and subject to dynamic exclusion for 30 s afterwards. Selected ions were sequentially subjected to collision-induced association (CID) activation conditions in the linear ion trap where sequence spectra were collected very quickly. The AGC target for CID was 4.0E04, 35 % normalized collision energy (NCE), with an activation Q of 0.25 and a 75 ms maximum fill time. Identical precursors were subjected to a 2nd MS2 also performed in the trap, but the top 10 sequence ions were selectively retained. These are routed backward to flow ‘upstream’ to the High energy Collision Dissociation (HCD) cell and activated at 60 % NCE to dissociate smaller reporter ions. HCD fragment ions were analyzed using the Orbitrap (AGC 1.2E05, with a resolution set to 50,000 and a mass range restricted to 80-500 m/z).

Initial spectral searches were performed with ProteomeDiscoverer3.0.1 (Thermo, PD3) against a UniProtKB mouse database. An equal number of decoy entries were created and searched simultaneously by reversing the original entries in the target databases. Precursor mass tolerance was set to 10 ppm and fragments acquired in the Linear Trap were searched at 0.4 Da and reporters (MS3) were acquired at a resolution of 50,000. Data files were loaded into Scaffold Proteomics Software for quantitative analysis across groups. To differentiate between the effects of aging and the effects of exercise during aging, we compared eAMD vs. HYA and eAMD+Ex vs. eAMD separately. To fully characterize our models, we considered all proteins that were significantly altered, regardless of log_2_ fold change cut-offs. Further statistical analyses can be seen below.

### Bioinformatic analyses

Several tools were used to characterize the proteome of isolated MAMs. MetaMass, a publicly available template for excel and R, comprises a comprehensive list of putative subcellular markers gathered from numerous publications and databases such as UniProt Knowledgebase (UniProtKB) and Human Protein Atlas (HPA) (Lund-Johansen et al, 2016). Using this, we compared our proteome to others that included proteins only annotated to a single subcellular compartment. Overlap of proteins (or lack thereof) was detected using the conditional formatting function within Microsoft Excel. The clusterProfiler package in R was used to perform Gene Set Enrichment Analysis of altered GO terms in the two group comparisons (eAMD/HYA and eAMD+Ex/eAMD) (Wu et al, 2021). ClusterProfiler graphs were downloaded as scalable vector graphics (SVG) and visually adapted using the open-source, vector graphics editor InkScape®. All diagrams were created using BioRender Illustration Software.

### Statistical analysis

One-way ANOVA analysis was used to compare HYA, eAMD, eAMD+Ex or HYA, eAMD and aAMD groups. The Kruskal-Wallis test was performed to analyze differences in the number of contact sites per mitochondrion across HYA, eAMD, and eAMD+Ex groups. Data are presented as mean ± SE. Violin plots are represented with first quartile, median, and third quartile. Proteomic comparisons were conducted using a Mann-Whitney U Test. Values of p < 0.05 were considered significant. All statistical tests were performed using GraphPad Prism software v. 10.3.1 (GraphPad Software, Boston, MA, USA).

## Results

### Regular exercise prevents morphological and functional aspects of early dysfunction in aging muscle

To elucidate mechanisms preceding the onset of advanced age-related muscle dysfunction (aAMD), we first compared animals of 6, 21 and 31 months of age. Compared to healthy young adults (HYA), animals at 31 months of age displayed significantly lower hindlimb muscle mass (i.e., of GA, PL and TA) and force (Figure S1). Animals at 21 months of age had unaltered TA and PL muscle masses, a modest decline in GA muscle mass, and preserved muscle force (Figure S1). Based on these data, and a recently developed definition of sarcopenia in mice consistent with the European Working Group on Sarcopenia in Older People (EWGSOP) (i.e., low muscle strength, muscle mass and physical performance in relation to young adult counterparts) (Kerr et al, 2024), we considered this group to be at an early stage of age-related muscle dysfunction (eAMD) (Figure 1a). Next, we investigated the impact of regular exercise during eAMD by subjecting a separate cohort of 19-month old animals to 6-8 weeks of progressive treadmill training (i.e., finishing at 21-months of age; eAMD+Ex group). This exercise regimen was sufficient to prevent bodyweight gain associated with sedentary aging and improve exercise capacity (Figure 1b and Figure S2A,B). Moreover, regular exercise effectively prevented age-related atrophy in GA muscles, increased plantaris (PL) muscle mass, without altering TA muscle mass (Figure 1c-e). Of note, PL fiber type composition was not altered in eAMD animals indicating that the denervation and neuromuscular junction impairments that preferentially affect MyHC 2X and 2B fibers at more advanced stages of age-related dysfunction were not yet present (Figure 1f,g). The exercised group had a larger overall fiber size driven by MyHC 2A, 2X and hybrid fibers (Figure 1f, g, Figure S3). Neither age nor exercise altered *in vivo* maximal or submaximal torque (Figure 1h). However, at a submaximal stimulation frequency, eAMD animals were approximately 10 % closer to attaining their maximal torque value(s) than HYA, and this was prevented by regular exercise (Figure 1i). Without changes to fiber composition, the attainment of a higher relative force at a submaximal stimulation frequency may result from slower ECR cycles (Mayfield et al, 2023). Accordingly, eAMD animals did not present deficits in the rate of force development but exhibited reductions in both maximum and mean rate of relaxation compared to HYA. Exercise also reverted this outcome (Figure 1j-l). eAMD animals were more prone to peripheral fatigue than young and exercise-trained animals, as indicated by a larger decrease in force (58 % vs. 43 % and 48 %, respectively) during repeated submaximal contractions (Figure 1m). As expected, all groups presented a progressive deterioration in the rate of relaxation during the fatigue protocol, but such decline was more pronounced in eAMD animals (Fig 1n). Collectively, these data highlight the slowing of muscle relaxation as a critical functional deficit occurring early in the development of age-related muscle dysfunction, and that regular endurance exercise prevents this outcome.

**Figure 1.**
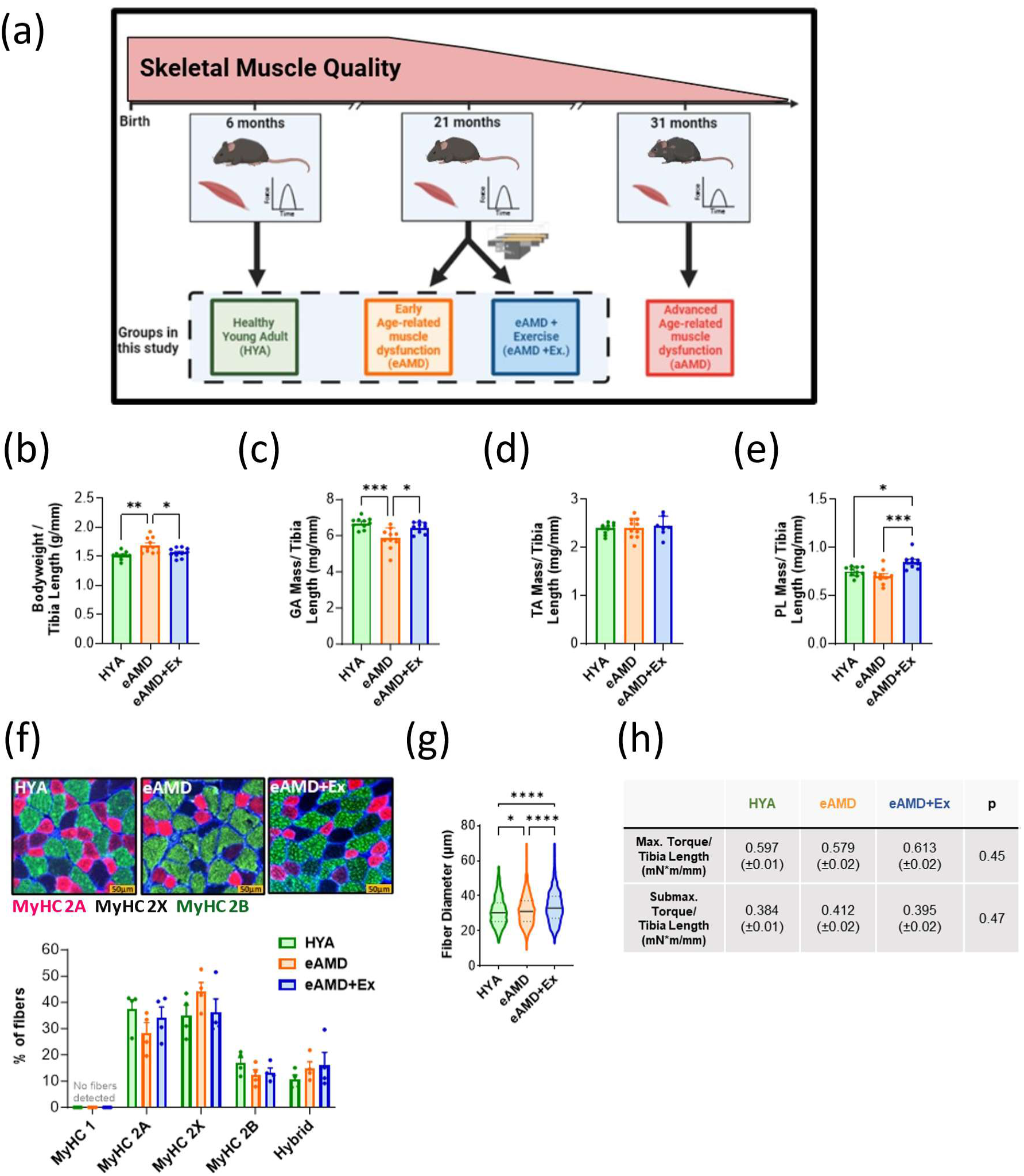
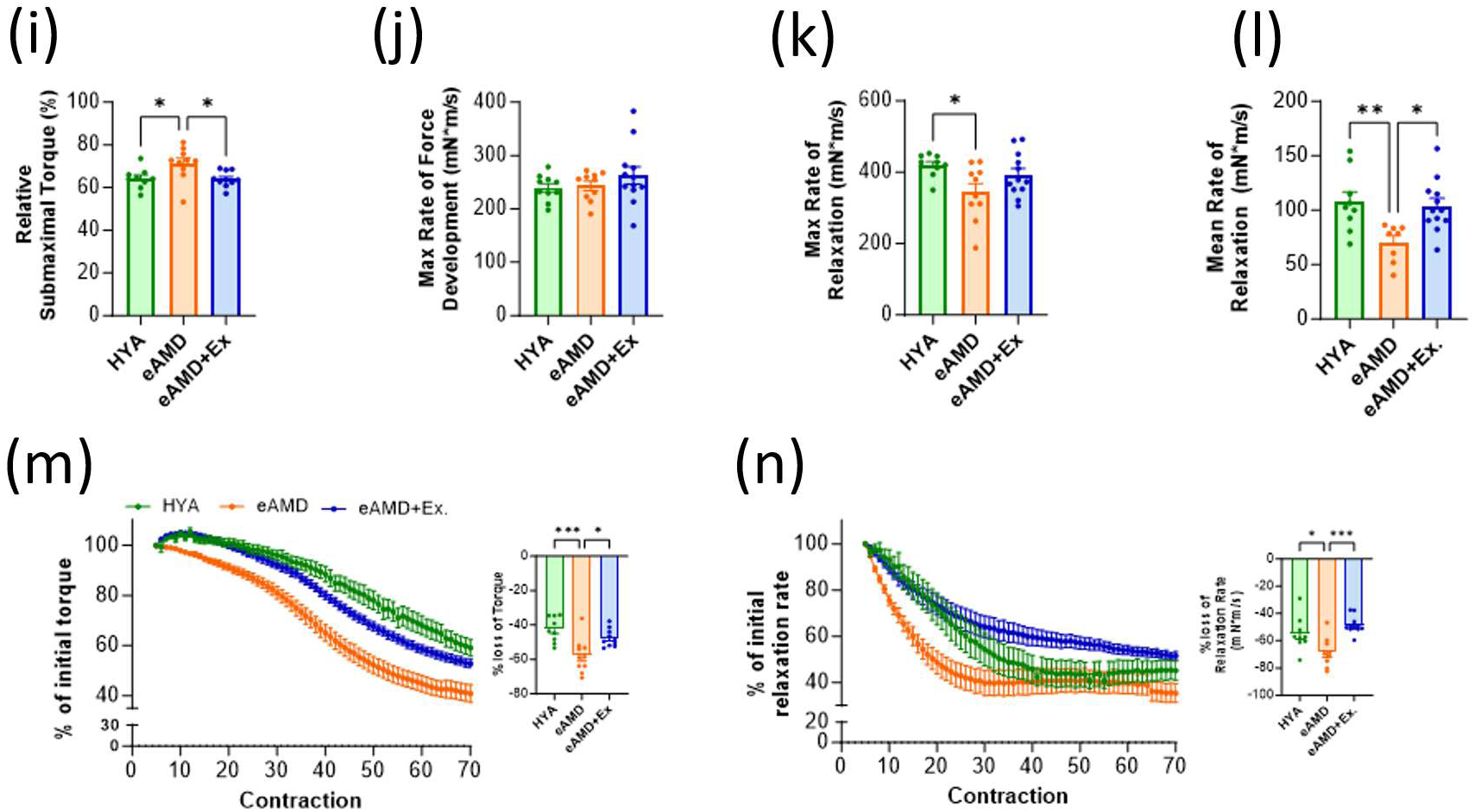
Effects of aging and exercise on skeletal muscle characteristics. (a) Diagram depicting skeletal muscle health across the lifespan and groups involved in the present study. (b) Bodyweight and wet muscle weight for (c) gastrocnemius, (d) tibialis anterior, and (e) plantaris wet weight normalized to tibia length (n=10). (f) Top-Representative images of immunofluorescent (IF) staining of MyHC isoforms in each group. Bottom-Percentage of different MyHC isoforms in PL muscle fibers (n=4). (g) PL fiber diameter including all fiber types. Violin plots display 1st quartile, median, and 3rd quartile. (h) Summary table of maximal (150Hz) and submaximal (50Hz) isometric torque of plantar flexors normalized to tibia length (n=10-12). (i) Submaximal torque expressed as a percentage of maximal torque. (j) Maximal rate of force development during 150Hz stimulation. (k) Maximal rate of relaxation during 150Hz stimulation. (l) Average rate of relaxation observed during 70-repetition stimulation protocol. (m) Left-torque values at each contraction averaged within groups during repetitive stimulation protocol. Right-percentage of initial torque lost after 70 contractions at 50Hz (n=9-12). (n) Left-rates of relaxation at each contraction averaged within groups during repetitive stimulation protocol. Right-percentage of initial relaxation rate after 70 contractions at 50Hz. Data are means ± SEM; *p<0.05, **p<0.01, ***p<0.001, ****p<0.0001. HYA – Healthy Young Adult, eAMD – early Age-related Muscle Dysfunction, eAMD+Ex – early Age-related Muscle Dysfunction following 6-8 weeks of regular endurance exercise.

### Exercise prevents early age-related changes in mitochondrial ultrastructure and ROS emission

The observed deficits in muscle relaxation suggested that bioenergetics may be impaired early with aging. Therefore, we next assessed mitochondrial ultrastructure in red gastrocnemius (GA) fibers using transmission electron microscopy (TEM) imaging to identify potential intrinsic mitochondrial deficits in eAMD. We observed lower surface area and circularity of individual intermyofibrillar (IM) mitochondria in eAMD (Figure 2a-c). Exercise attenuated the decrease in surface area of individual mitochondria while increasing their circularity above what was observed in HYA muscle (Figure 2a-c). These ultrastructural changes did not seem to translate into differences in mitochondria content across groups, since relative mitochondrial area in TEM images remained unchanged and the expression of electron transport chain (ETC) subunits was not consistently higher in eAMD+Ex vs. eAMD groups (Figure 2d, Figure S4A).

**Figure 2.**
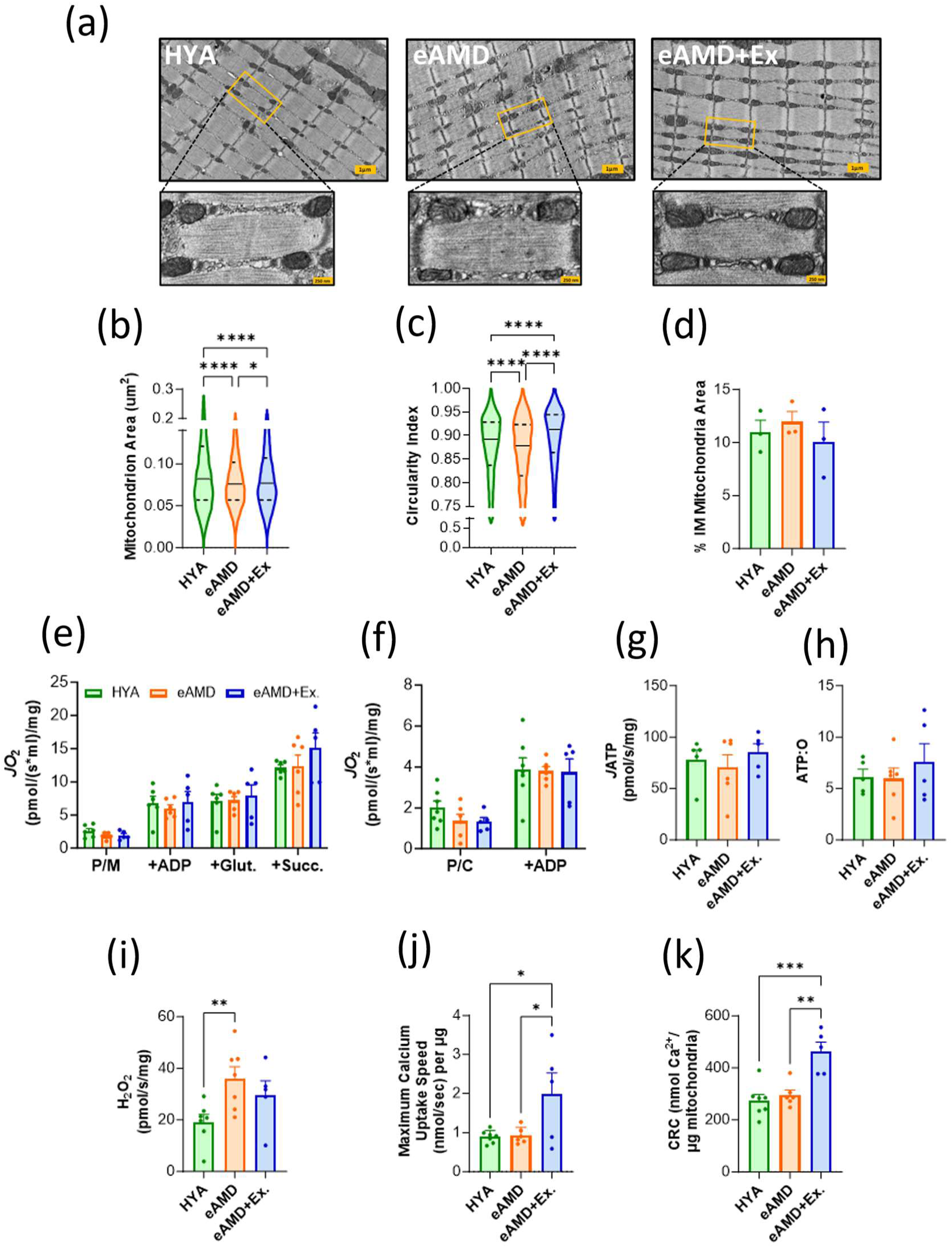
Effects of aging and exercise on skeletal muscle mitochondria. (a) Representative TEM images of intermyofibrillar (IM) mitochondria in red gastrocnemius from each group (n=3 animals, 3-4 images per animal). (b) Average surface area of individual IM mitochondria. (c) Circularity index (Y= 4π *[SA/(perimeter2)] of individual mitochondria. (d) Percentage of quantified area occupied by IM mitochondria. (e) High-resolution respirometry measurements of oxygen flux (*J*O_2_) using carbohydrate-based and (f) fatty acid-based substrates in permeabilized red gastrocnemius fibers. (g) ATP production rate (*J*ATP) in permeabilized fibers after injection of ADP; derived via NADPH autofluorescence. (h) Total ATP production from (g) normalized to the amount of oxygen consumed in the same amount of time. (i) Hydrogen peroxide (H_2_O_2_) emission following injection of succinate (no ATP) in permeabilized red gastrocnemius fibers (n=5-7). (j) Highest rate of Ca²⁺ uptake achieved during a single Ca²⁺ pulse in isolated TA mitochondria (n= 5-7). (k) Total Ca^2+^ retention capacity (CRC) prior to swelling and mitochondrial permeability pore (mPTP) opening. P/M-Pyruvate/Malate; Glut - Glutamate; Succ-Succinate; P/C-Palmitoylcarnitine. Data are means ± SEM; Violin plots display 1st quartile, median, and 3rd quartile. Bar graphs are means ± SEM; *p<0.05, **p<0.01, ***p<0.001., ****p<0.0001. HYA – Healthy young adult, eAMD – early Age-related Muscle Dysfunction, eAMD+Ex – early Age-related Muscle Dysfunction following 6-8 weeks of regular endurance exercise.

We then interrogated mitochondrial respiratory function in saponin-permeabilized red gastrocnemius (GA) fibers. Contrary to expectation, maximal rates of respiration (*J*O_2_) utilizing either carbohydrate-based or fatty acid-based substrates were not different across groups (Figure 2e,f). The efficiency of ATP synthesis, measured by the P:O ratio and rate of ATP production (*J*ATP), was also not affected in early age-related muscle dysfunction or by regular exercise (Figure 2g,h). However, fibers from eAMD muscles exhibited higher levels of H_2_O_2_ emission compared to HYA, and this was mitigated by exercise (Figure 2i). Even though relative GSH:GSSG ratios did not reach statistical significance across groups, eAMD muscles had a ∼50% lower ratio than the other groups suggesting an increased oxidant load (Figure S4B-D).

Because mitochondrial Ca^2+^ uptake directly regulates various TCA cycle enzymes and ROS production, we then evaluated Ca^2+^ uptake by isolated mitochondria from the tibialis anterior (TA) muscle. Neither maximal Ca^2+^ uptake rate nor Ca^2+^ retention capacity was altered in eAMD (Figure 2j, k). However, exercise significantly increased both measures of mitochondrial calcium handling (Figure 2j, k). Overall, these results highlight subtle abnormalities in intermyofibrillar mitochondria ultrastructure and increased ROS emission as potential drivers of the phenotype in eAMD despite preserved overall mitochondrial content, Ca^2+^ uptake dynamics and maximal respiratory capacity. Importantly, regular exercise effectively prevented these early aging-related changes.

### MERC ultrastructure is altered with age and normalized by exercise

Since mitochondrial respiratory function was preserved in early age-related muscle dysfunction and unaffected by exercise, we then hypothesized that MERC ultrastructure would be altered at that stage potentially contributing to slower relaxation rates by compromising the communication between mitochondria and the SR. High-magnification TEM images (8000X) of the red portion of GAs were captured to assess MERC ultrastructure using previously described parameters (Figure 3a) (Giacomello et al, 2016). MERC width describes the cytosolic cleft distance between the SR membrane and the mitochondrial outer membrane. Because many of the molecules exchanged at MERCs are subject to the diffusion laws of Einstein and Fick, a larger separation (i.e., increased width) between mitochondria and SR would decrease the efficiency of nucleotide (e.g., ATP) and/or ion (e.g., Ca^2+^) transport across organelles (Giacomello et al, 2016). On the other hand, MERC length defines the stretch of individual SR membrane segments parallel to the outer mitochondrial membrane at contact sites. Compared to HYA, MERC width was not altered in eAMD or eAMD+Ex MERCs. This suggests that exchange efficiency was preserved at this early stage of muscle dysfunction (Figure 3b).

**Figure 3.**
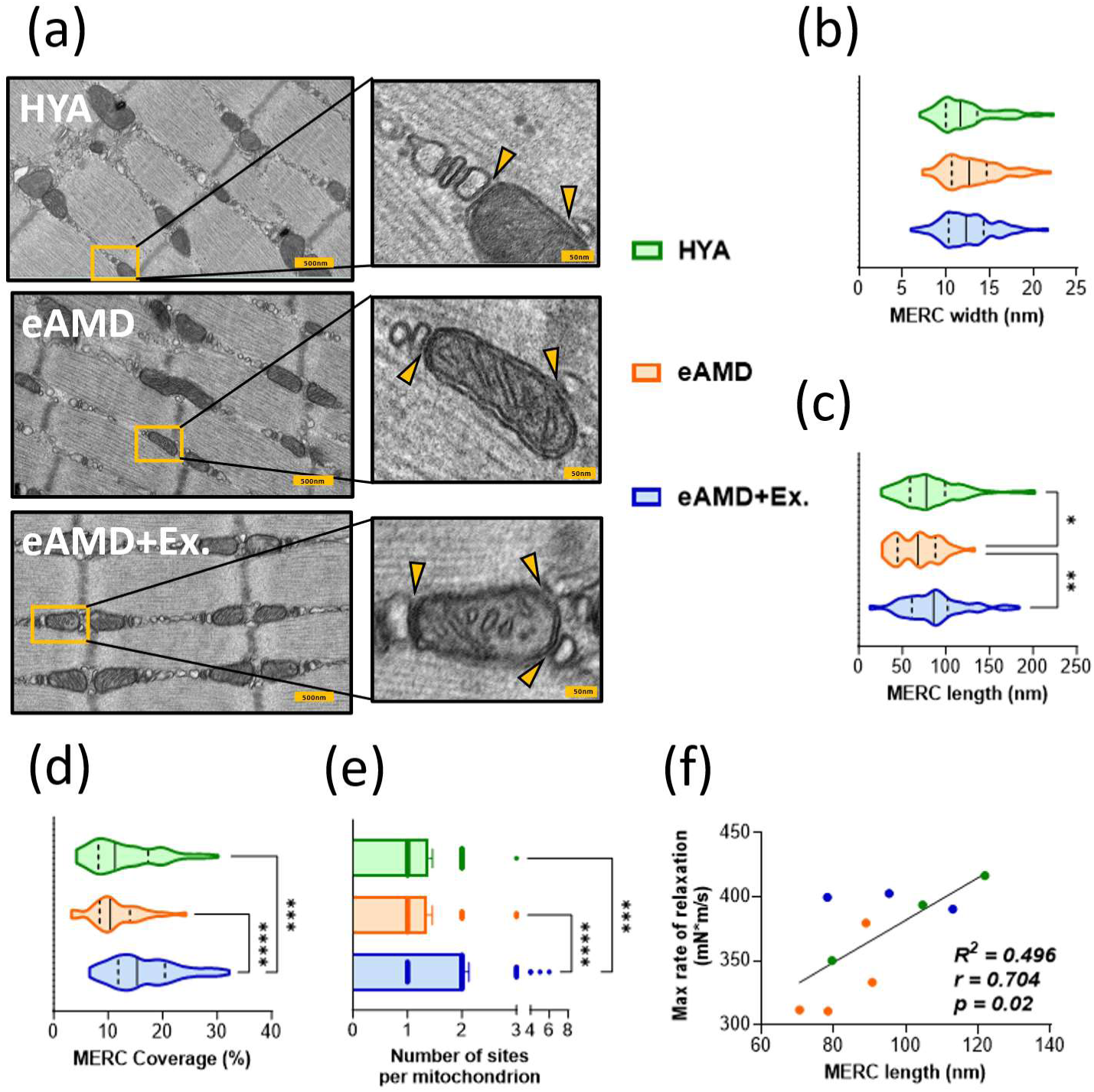
Impact of aging and exercise on skeletal muscle MERC ultrastructure. (a) Representative TEM images of quantification of MERCs in red gastrocnemius. Yellow arrows indicate MERC region of interest (i.e., physical contact between mitochondria [gray] and SR [white]) (n=3 animals/group, 2-3 images per animal, 5-10 sites per image). (b) Distance between the cytosolic face of sarcoplasmic reticulum (SR) and the outer mitochondrial membrane (OMM). (c) Length of individual SR membranes in contact with OMM. (d) Relative percentage of mitochondrion perimeter opposed by SR membrane contact(s). (e) Number of ER/SR contact sites per mitochondrion (MERCs) in each group. (f) Pearson correlation coefficient (*r*), and coefficient of determination (*R^2^*) resulting from MERC length and Maximal rate of muscle relaxation across the three groups (n=3/group). Violin plots display 1st quartile, median, and 3rd quartile. *p<0.05, **p<0.01, ***p<0.001., ****p<0.0001. HYA – Healthy young adult, eAMD – Early age-related muscle dysfunction, eAMD+Ex – Early age-related muscle dysfunction following 6-8 weeks of regular endurance exercise.

However, MERC length was significantly lower with aging, and this was prevented with regular exercise (Figure 3c). Assuming that MERC length is proportional to the cumulative transport potential of molecules across the two organelles, MERCs in eAMD may have a diminished capacity for exchanging nucleotides and ions between the SR and mitochondria potentially impacting contractile function. Notably, MERC length was strongly associated with the *in vivo* rate of muscle relaxation across groups, with the former explaining ∼50% of the variation of the latter (Figure 3f, Figure S5A-C). Along with the length of individual MERC sites, we also assessed cumulative MERC lengths relative to the respective mitochondrion perimeter (i.e., MERC coverage). MERC coverage was unaltered in early aging but was increased with regular exercise even when compared to healthy young adults (Figure 3d). This trend may be partially attributable to a greater number of SR tubules per mitochondrion in animals that exercised (Figure 3e). These data suggest that MERC ultrastructure is altered in early AMD and that the exercise modulation of MERC ultrastructure is beneficial to muscle function.

### MAM protein composition is differently impacted in early aging and by regular exercise

Since MERC ultrastructure was altered with aging and exercise, we next interrogated if changes in proteins expressed at MERCs would be associated with early age-related muscle dysfunction and the benefits of exercise. To address this question, we first isolated mitochondria-associated endoplasmic reticulum membranes (MAMs) following a previously established protocol (Wieckowski et al, 2009) with a few modifications (see Materials and Methods section), and then probed equal amounts of fractionated proteins for markers of specific cellular compartments. The cytosolic protein α-tubulin was present in the whole tissue lysate and in the ER/SR fraction. Conversely, the residing ER/SR protein Eukaryotic Translation Initiation Factor 2 Alpha Kinase 3 (EIF2K3, also referred to as PERK) was enriched in the ER/SR fraction. Both α-tubulin and PERK were undetectable in the crude mitochondria and MAM fractions. In this protocol, MAM fractions are obtained from the crude mitochondria fraction. Therefore, while MAM fractions contained only traces of the mitochondrial matrix protein Citrate Synthase (CS), both fractions were enriched with the canonical MAM marker, Acyl-CoA Synthetase Long Chain Family Member 4 (ACSL4) (Figure 4a) (Wieckowski et al, 2009). Of note, total protein content in MAM fractions relative to the amount of GA tissue processed did not differ across groups indicating that overall quantitative changes in the MAM proteome are not present at this early stage of aging (Figure 4b). Next, we TMT-labeled MAM fractions from the three groups and subsequently submitted those samples through LC-MS identifying a total of 473 proteins. To gain insight into the cellular locations of these proteins, we indexed our MAM proteome across the Human Protein Atlas (HPA), UnitProt/GO using MetaMass (Lund-Johansen et al, 2016). Accordingly, ∼75 % of the proteins in these fractions were located at either the mitochondrion or ER/SR. The next most abundant cellular compartment for the identified proteins was the cytosol. This was not unexpected as some MERC proteins do not physically interact with the ER/SR or the outer mitochondrial membrane but are located between these organelles (Figure 4c) (Zhou et al, 2023). Subsequently, we juxtaposed our proteome to previous investigations of MAM fractions. Over 85 % of identified proteins (i.e., 417 proteins) aligned with work recently performed in rat skeletal muscle, indicating that the protein composition of skeletal muscle MAMs is well-conserved in rodents (Supplementary Document 1) (Lu et al, 2022). Among those, 105 proteins aligned with previously established consensus MAM proteins, which were based on studies of MERCs across a myriad of non-contractile cells and tissues (Carreras-Sureda et al, 2019). The remaining group of 312 MAM proteins appears unique to skeletal muscle given their presence in muscle MAM fractions obtained from mice (current work) and rats (Lu et al, 2022) (Figure 4d).

**Figure 4.**
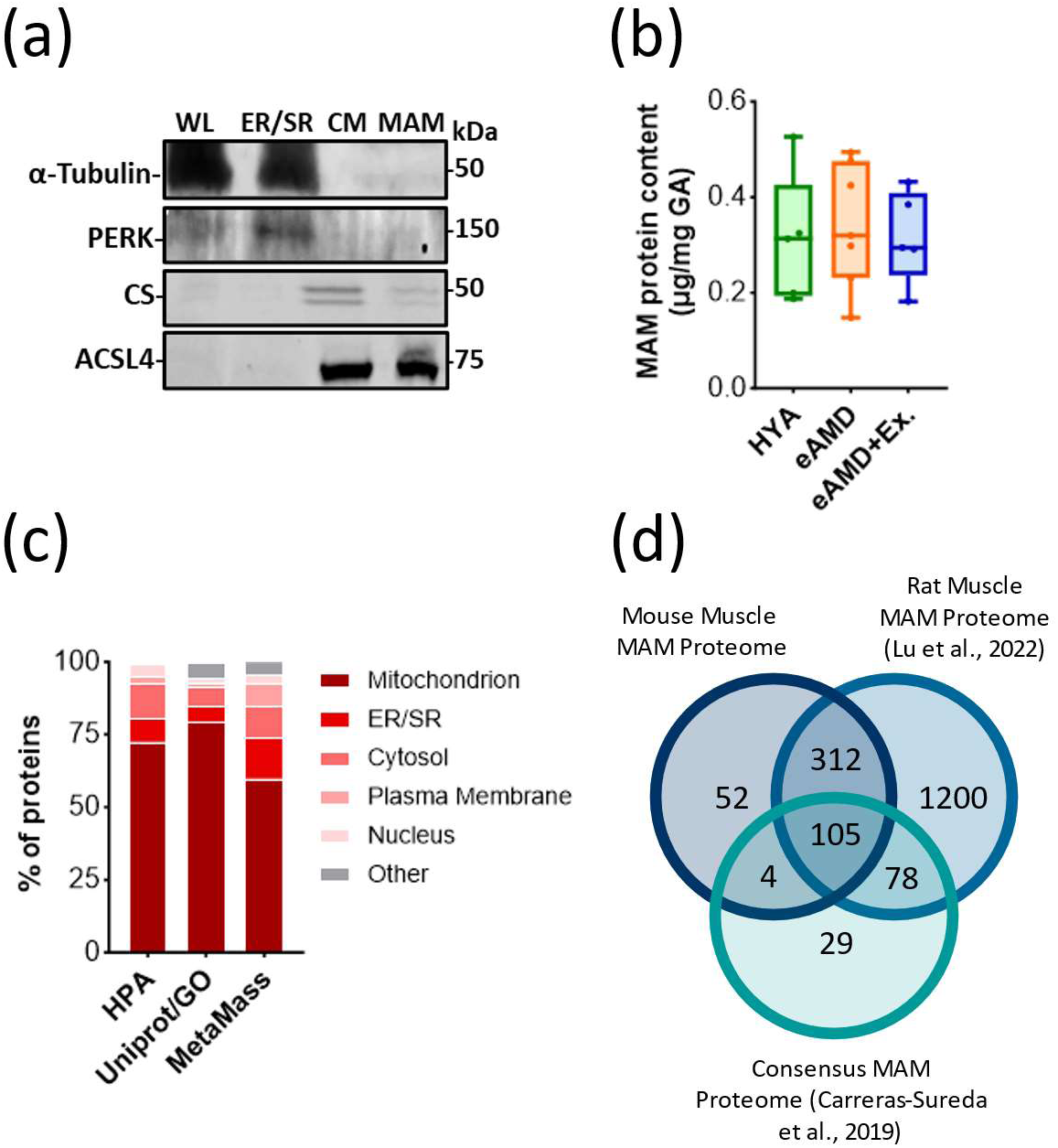
Characterization of skeletal muscle MAM proteome. (a) Representative immunoblots of muscle tissue fractions denoting the obtained fractions of whole lysate (WL), endoplasmic and sarcoplasmic reticulum (ER/SR), crude mitochondria (CM) and MAMs. (b) Total protein yield of MAM isolations normalized to amount of GA tissue used (n=3-4). (c) Distribution of proteins with single-annotation in the MAM proteome according to HPA, UniProt and MetaMass. (d) Comparisons between mouse skeletal muscle MAM proteome (current study), rat skeletal muscle MAM proteome (Lu et al., 2022), and consensus MAM proteins (Carreras-Sureda et al., 2019). HYA – Healthy young adult, eAMD – Early age-related muscle dysfunction, eAMD+Ex – Early age-related muscle dysfunction following 6-8 weeks of regular endurance exercise.

To interrogate the potential effects of early biological aging on skeletal muscle MAM protein composition, we then compared the subcellular proteomes of HYA and eAMD animals. Out of the 473 MAM proteins identified, 121 changed with age. From those, 73 proteins (15 %) were increased, and 48 proteins (10 %) decreased in eAMD (Figure 5a, Supplementary Document 2). Due to the dynamic localization of proteins at MAMs, we sought to characterize which subcellular locations were most affected. To this end, we analyzed the 121 significantly-altered MAM proteins using Gene Set Enrichment Analysis (GSEA) via Gene Ontology. In terms of cellular compartments (i.e., GO:CC), eAMD MAMs displayed an increased enrichment of SR/ER-related proteins, while HYA MAMs were mostly enriched with Mitochondrial Envelope proteins (Figure 5b, Supplementary Document 3). Related to biological processes (i.e., GO:BP), proteins involved in the removal of ROS (e.g., cytochrome c (Cycs), superoxide dismutase 2 (SOD2), and nicotinamide nucleotide transhydrogenase (Nnt)) and regulation of membrane potential were enriched in HYA MAMs, while eAMD MAM fractions displayed enrichment of proteins responsive to glycolytic and catabolic substrates (Figure 5c; Supplementary document 4).

**Figure 5.**
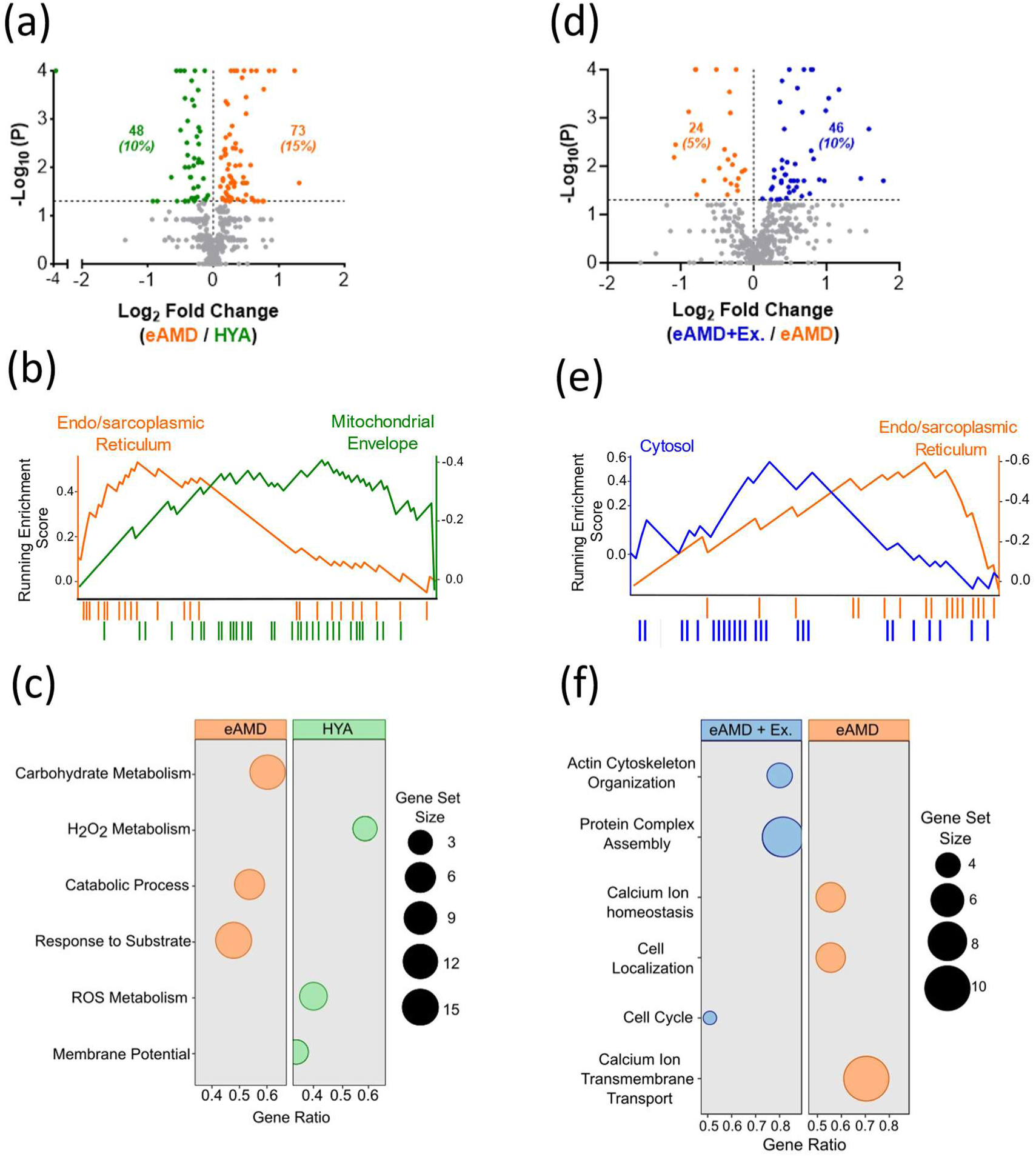
Effects of aging and exercise on skeletal muscle MAM proteome. (a) Volcano plot comparing the MAM proteome of eAMD vs. HYA (n=3). Number and percentage of differentially expressed proteins (colored) relative to the entire proteome (473 proteins) are shown. Horizontal dashed line denotes the lower limit for significance (i.e., -log10(p) >1.3). (b) Top enriched (padj < 0.05) discrete organellar proteins (genesets) from GO:CC (GSE analysis) of differentially expressed genes or proteins (DEGs) from eAMD vs. HYA comparison in ClusterProfiler R package (Outcome was N = 25 and 33 total proteins for endo/sarcoplasmic reticulum and mitochondrial envelope, respectively). (c) Top enriched genesets (padj < 0.05) from GO:BP (GSE analysis) in ClusterProfiler R package. (d) Volcano plot comparing proteome of eAMD+Ex vs. eAMD MAMs (n=3). (e) Top enriched (padj < 0.05) discrete organellar proteins (genesets) from GO:CC of DEGs from eAMD+Ex vs. eAMD comparison. (f) Top enriched genesets (padj < 0.05) from GO:BP in eAMD+Ex vs. eAMD.

To examine if changes in the MAM proteome underlie the beneficial adaptations of muscle to regular exercise, we compared eAMD and eAMD+Ex groups. Of the 473 proteins identified, 70 significantly changed with exercise. Of those, 46 proteins (10 %) were increased in eAMD+Ex, while 24 proteins (5 %) were decreased (Figure 5d, Supplementary document 2). In terms of cellular compartments, eAMD MAMs again displayed enrichment of ER/SR-related proteins when compared to eAMD+Ex. Interestingly, the top-enriched, discrete cellular compartment for proteins changing with exercise was the cytosol, suggesting exercise-specific adaptations to MAMs (Figure 5e, Supplementary Document 5). The biological processes mostly affected by regular exercise comprised broad categories, including cytosolic modifications (e.g., actin cytoskeleton organization and protein complex assembly) and cell cycle (Figure 5f). Comparatively, the pool of MAM proteins enriched in aging sedentary muscle suggested an increased reliance on ER/SR-regulated processes related to calcium homeostasis (e.g., calcium ion homeostasis and transmembrane transport) (Figure 5f, Supplementary Document 6). Altogether, these data support a paradigm in which exercise not only prevents changes in ER/SR proteins at MAMs observed early in sedentary aging, but also promotes the enrichment of cytosolic proteins at MAMs likely contributing to improvements in ER/SR-mitochondria cross-talk.

### Aging and exercise inversely modulate the expression of several MAM proteins

To identify MAM proteins most relevant to muscle health during aging, we collectively examined proteins that were inversely modulated by sedentary aging and exercise. In total, 28 proteins fell into this category. Of those, 17 were higher, while 11 were lower in HYA and eAMD+EX vs. eAMD (Figure 6a). Interestingly, all proteins present at higher levels in MAMs of young adult and exercised animals were located in either the mitochondrion or the cytosol and were generally involved in energetics and cytoskeleton integrity. On the other hand, most of the proteins present at higher levels in MAMs of animals with early age-related muscle dysfunction were located in the ER/SR and were generally involved in protein quality control and calcium dynamics (Figure 6a,b, Supplementary document 7). We next tried to obtain insights into whether those changes were secondary to the expression of these proteins in the whole muscle tissue. To address this, we assessed the expression of MAM proteins found at the SR and at the mitochondrial outer membrane (i.e., ATP2A1 and VDAC1, respectively) in whole GA muscle lysates. Their whole tissue expression was unaltered with sedentary aging or with exercise, despite their either increased (i.e., ATP2A1) or decreased presence at MAMs (i.e., VDAC1) of eAMD vs. HYA and eAMD+Ex (Figure 6a,c,d). Collectively, these observations suggest that at least some of the MERC protein changes occurring in early age-related muscle dysfunction result from their altered localization at the contact sites and that regular exercise can prevent this.

**Figure 6.**
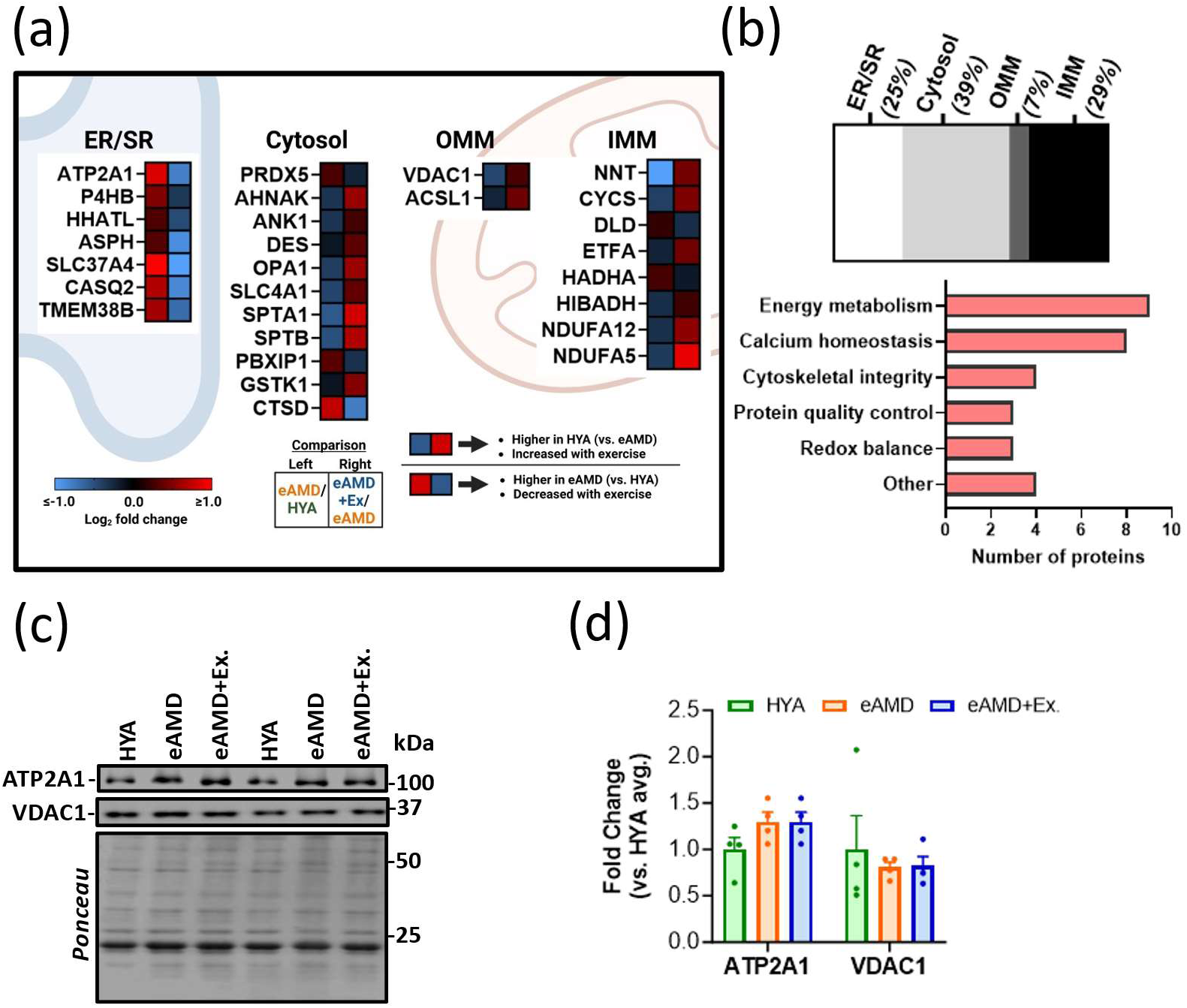
MAM proteins inversely modulated in early sedentary aging and regular exercise. (a) Inversely regulated MAM proteins (with age and exercise) according to their location at MERCs. Heatmaps represent respective quantitative proteomic comparisons (refer to keys at the bottom). (b) Distribution of inversely modulated MAM proteins across MERC subcellular locations (top) and molecular functions (bottom). These were collectively curated from NCBI and GeneCards. (c) Representative immunoblots and (d) Quantification of ATP2A1 and VDAC1 expression in whole GA lysates (n=4). HYA – Healthy young adult, eAMD – Early age-related muscle dysfunction, eAMD+Ex – Early age-related muscle dysfunction following 6-8 weeks of regular endurance exercise.

## Discussion

Due to their terminal differentiation, skeletal muscle fibers are particularly susceptible to biological aging. After age 50 in humans, muscle mass decreases at ∼1-2 % per year and this rate nearly doubles after 60 (von Haehling et al, 2010). Occurring in parallel is a decrease in force output that reduces physical function, compromises independence, and increases the risk of mortality (Lang et al, 2010). Therefore, the most effective treatments for sarcopenia are likely those that are targeted at the earliest stages of aging. Regular exercise is currently the only effective therapy for age-related muscle dysfunction. However, many social, economic, and health-related barriers exist that prevent individuals from maintaining consistent activity. Despite our considerable understanding of the maladaptive processes transpiring at advanced stages of muscle aging, our knowledge of the initial molecular events leading to sarcopenia remains elusive. A better understanding of this is therefore required to begin developing effective therapies. Based on a recently adapted definition from EWGSOP for sarcopenia in mice, we studied a group of animals considered to be at an “early” stage in the progression of sarcopenia or age-related muscle dysfunction (eAMD) (Kerr et al, 2024). These mice presented either mild or no muscle atrophy with preserved maximal force. Despite this, we show that relaxation was much slower in these mice. As a result, their E-C-R cycles were elongated, creating premature tetanic fusion and higher force –relative to their max– during submaximal contractions. Though this may be partially beneficial for a few successive muscle contractions, we observed that it predisposes the muscles to fatigue over a longer period of time. This is supported by recent data in humans that showed an association between longer relaxation time and perceived fatigue (Mast et al, 2024). Since their fiber type distribution was unaltered, these functional changes took place before age-related denervation and subsequent type 2 to type 1 fiber switching. Of note, because our functional measurements involve direct stimulation of the tibial nerve, and because motor neurons conduct electrical signals (i.e., action potentials) ∼100 times faster than muscle fibers, we can rule out neuronal-mediated deficits as a cause of reduced muscle relaxation (Loeb et al, 2000; Zollinger et al, 2015). Collectively, these observations indicate that the slowing of muscle relaxation is likely the first functional impairment to occur with aging, and that its driving mechanisms reside within the myofibers.

To begin interrogating potential causes of this, we studied an additional group of eAMD mice that regularly exercised. Our 8-week intervention was effective at preventing the decline of relaxation rate while also mitigating muscle fatigue. Due to the heavy reliance of these contractile functions on a constant supply of ATP, we turned our attention to muscle mitochondria. In fact, dysfunction of mitochondria occurs ubiquitously in more advanced stages of aging (Marzetti et al, 2024). Albeit intermyofibrillar mitochondria content was unchanged in our model, those were smaller and less circular in eAMD muscles. In a two-dimensional sense, this equates to a decreased surface area to circumference ratio, which may result in a lowered ability of mitochondria to exchange material with other organelles. Interestingly, maximal respiratory capacity was largely unaffected by aging and exercise, suggesting that the bioenergetic capacity of mitochondria is preserved at early stages of sarcopenia. Mitochondrial H_2_O_2_ emission, however, was increased with aging and attenuated by exercise. These findings implicate elevated mitochondrial ROS emission as an important (mal)adaptation in muscle that precedes the major contractile deficits normally observed during advanced aging. Because the oxidation of macromolecules has been shown to compromise muscle function in different settings, future studies are required to clarify a) which specific molecules may be modified by mitochondrial ROS emission and b) whether these are related to alterations in muscle relaxation (Andersson et al, 2011; Powers et al, 2011).

Within muscle fibers, Ca²⁺ is not only critical for contractile function, but it also stimulates oxidative phosphorylation when imported into mitochondria (Duchen, 2000). In that sense, an impaired ability of mitochondria to uptake Ca²⁺ could limit the acceleration of ATP resynthesis required for SERCA-mediated SR Ca²⁺ reuptake, thus delaying muscle relaxation. However, our results demonstrated that when exposed to excess Ca²⁺, neither speed of mitochondria Ca²⁺ uptake nor total retention capacity were impacted in early aging muscle. Though exercise improved these to levels above HYA animals, the previous observation suggests that the intrinsic capacity of mitochondria to import Ca²⁺ is not a main culprit in the slowing of relaxation seen in eAMD. Next, we asked whether external mitochondrial interactions could be responsible for the phenotype observed in early aging. Indeed, such interactions are becoming increasingly recognized as critical regulators of intra and inter-organelle homeostasis (Elbaz-Alon et al, 2017). In skeletal muscle, intermyofibrillar MERCs are particularly important as they represent physical interactions between the critical site of Ca^2+^ handling and regulation of contraction (i.e., the ER/SR) and ATP resynthesis (i.e., mitochondria). Therefore, we first examined these MERCs using well-established morphological parameters: MERC width and MERC length (Giacomello et al, 2016). According to laws of diffusion, MERC width is thought to alter the exchange efficiency of metabolites across organelles (Giacomello et al, 2016). Our findings indicated that MERC exchange efficiency was unaltered in eAMD and regular exercise groups, and likely did not contribute to the contrasting muscle phenotypes observed in these conditions. Interestingly, recent studies suggest that MERC clefts become wider in advanced sarcopenia, which may contribute to the exacerbated contractile dysfunction present at that timepoint (Lu et al, 2022). MERC length was lower in eAMD mice and restored with exercise. Assuming there is relation between MERC length and the number of structures capable of exchange, the former should define the exchange capacity of an individual contact site. Because Ca²⁺ released from the SR can stimulate mitochondrial metabolism and ATP exported from mitochondria is important for optimal SR ATPase (i.e., SERCA) function, a decreased capacity to exchange ions and nucleotides between the two organelles could directly lead to reductions in the rate of muscle relaxation. In fact, MERC length was able to explain ∼50% of relaxation rate across HYA, eAMD, and eAMD+Ex groups (Figure 3f). Overall, these observations highlight the potential impact of MERC ultrastructure on muscle contractile function. It is important to note that future studies using 3D reconstructions of muscle MERCs are required to reveal additional nuances related to changes at the contact sites during aging and/or exercise.

In order to execute their numerous cellular processes, MERCs rely on an eclectic assortment of localized proteins. Related to ECR cycles in muscle, it is likely that some of these are either directly (e.g., exchange of ions and nucleotides) or indirectly (e.g., redox balance) involved. We therefore sought to profile the MERC proteome across our three groups by isolating MAM-enriched fractions from muscle. Total MAM protein concentration, which was shown to decrease at more advanced stages of aging, did not differ across groups (Lu et al, 2022). Next, LC/MS of the TMT-labeled MAM fractions allowed us to determine both quantitative and qualitative changes in the proteome. As expected, 80-90% of the 473 discovered proteins were annotated to either the mitochondrion, ER/SR or the cytosol (Figure 4c). Proteins linked to ER/SR lumen, mitochondrial matrix, nuclear, and plasma membrane proteins were also present in the fractions. This could be at least partly due to physical interactions between these proteins and MERCs *in vivo*.

We observed that during early age-related muscle dysfunction, 25% of MAM proteins were altered, with 15% being upregulated and 10% downregulated compared to healthy young adult MAMs (Figure 5a). Regular exercise in the older animals changed 15% of MAM proteins, upregulating 10% and downregulating 5% (Figure 5d). GSEA of eAMD vs. HYA animals revealed that aging led to a greater enrichment of ER/SR-related proteins at MAMs (Figure 5b). On the other hand, HYA MAMs were enriched with mitochondrial proteins. This was an interesting finding, since “ER-predominant” MAMs have been reported at more advanced stages of sarcopenia and may be attributed to reduced muscle oxidative capacity (Lu et al, 2022; Migliavacca et al, 2019). Further analysis of GO:BP showed that whereas ER-predominant MAM proteins are involved in catabolic and/or glycolytic processes, mitochondria-predominant MAMs (i.e., HYA MAMs) primarily modulate redox-related processes (Figure 5c). Thus, the latter may be partially responsible for the differences in H_2_O_2_ emission we observed. Additional studies on muscle mitochondria energetics and redox balance, such as *in vivo* P-MRS, will be needed to provide further insights into this paradigm.

Compared to exercise-trained older animals, sedentary animals maintained a greater enrichment of ER-related proteins, further implicating ER-predominant MAMs in muscle dysfunction (Figure 5e). Specifically, these MAMs showed higher levels of proteins involved in Ca²⁺ homeostasis (e.g., ATP2A1/SERCA1, RYR1, and TRDN) (Supplementary document 5). This may be due to compensatory upregulations of these proteins in response to diminished Ca²⁺ release unit coupling that has been shown to occur with sedentary aging (Michelucci et al, 2021). We hypothesized that exercise would effectively revert ER-predominant MAMs to the mitochondria-predominant MAMs observed in HYA animals. Interestingly, MAMs from the eAMD+Ex. group instead displayed enrichment of cytosolic components (Figure 5e). Albeit some of the protein sets that were modified with exercise included mitochondrial proteins (e.g., NDUFA5/6/12), the majority were deputed to sarcoplasmic reorganization (e.g., SPTA1, SPTB, FLNC) (Supplementary document 5). In the context of MERCs, these may be relevant in restoring MERC stability and facilitating more efficient MERC morphology (e.g., greater MERC coverage).

Finally, we sought to identify proteins that are inversely modulated by aging and exercise. In this way, we could profile a specific subpopulation of MAM proteins relevant to muscle health. We identified 28 candidate proteins that fit this description. These included seven localized to the SR, eleven to the cytosol, two in the outer mitochondrial membrane (OMM), and eight in the inner mitochondrial membrane (Figure 6a). Further inspection of whole-tissue levels of markers for MERC sub-compartments (e.g., ATP2A1/SERCA1-SR, VDAC1-OMM) revealed that their overall expression was not changing in the myofibers (Figure 6c, d). Taken together with changes in the MAM proteome, this indicates that in HYA and eAMD+Ex., a larger proportion of VDAC1 and a lower proportion of ATP2A1 are localized to MERCs. Thus, the localization of proteins to the contact sites is regulated. Indeed, increased density of VDAC proteins to MERCs seems to allow microdomains of rapid Ca²⁺ transfer from the ER/SR to mitochondria, impacting the overall exchange between organelles (Rosencrans et al, 2021).

In conclusion, the present study provides foundational information regarding the impact of aging and regular exercise on skeletal muscle health, and MERC ultrastructure and protein composition. Our findings denote select atrophy of MyHC 2A fibers, preserved maximal torque, decreased relaxation rate and increased fatigability as features of early age-related muscle dysfunction that are reversed by regular exercise. Further, our data implicate mitochondrial ROS emission, and exonerate mitochondrial respiration, as a main subcellular culprit. For the first time, we reveal that the MAM ultrastructure and proteome are reciprocally modulated by aging and exercise. The identification of inversely regulated MAM proteins in with early age-related muscle dysfunction and exercise can be used to generate hypotheses investigating the influence of the contact sites and their protein composition on muscle dysfunction. Future studies focusing on the regulation and function of these proteins should provide new and important insights into the mechanisms modulating skeletal muscle health.

## Acknowledgments

We would like to acknowledge the University of Iowa Central Microscopy Research Facility and the University of Iowa Proteomics Facility, core resources supported by the University of Iowa’s Vice President for Research, and the Carver College of Medicine. We would also like to acknowledge Biorender.com, which was used in the creation of our diagrams. This work was supported by the NIH (R56AG068320), the Fraternal Order of Eagles Diabetes Research Center, the Office of the Vice-President of Research at the University of Iowa (V.A.L.). L.S was supported by NIH (HL130346, HL157741, and HL157781), R.O.P by NIH (DK125405), and E.J.A. by NIH (R01HL167087) and American Heart Association’s Strategically Focused Research Network award (20SFRN35200003).

## Author contributions

V.A.L. and R.O.P. conceived the original idea for this study. R.J.A., A.K., Q.S., M.P., E.C., and W.S. performed the experiments. L.S., E.J.A., and R.O.P. provided equipment and guidance for experiments. R.J.A analyzed the data and R.J.A., A.K., W.S., and V.A.L., interpreted the data. R.J.A., and V.A.L. wrote the manuscript. Q.S., M.P., L.S., E.J.A., and R.O.P. reviewed and revised the manuscript prior to submission.

## Data availability

Proteomics data and analysis (i.e., GO and GSEA) are available in supplementary documents. All other data may be supplied upon request from corresponding author.

---

**Supplementary figure 1.**
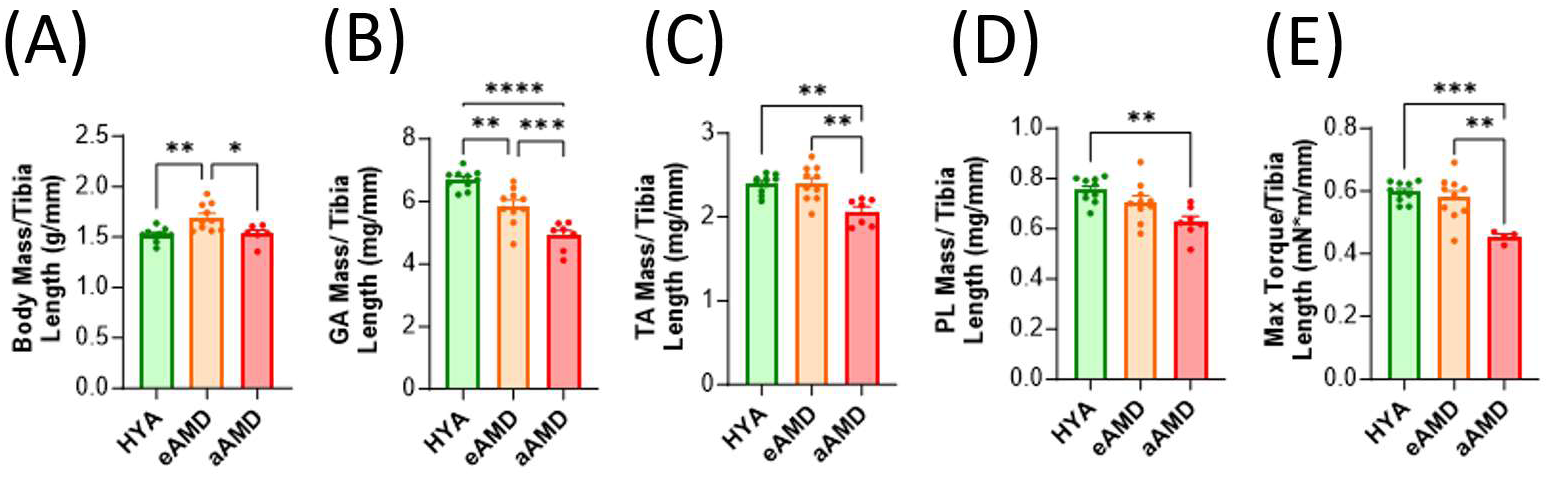
Phenotypic effects of aging in skeletal muscle. Refer to schematic in Figure 1A. (A) Bodyweight and wet muscle weight for (B) gastrocnemius, (C) tibialis anterior, and (D) plantaris muscles normalized to tibia length. (E) Maximal isometric torque of plantar flexors via stimulation (150 Hz) of the tibial nerve normalized to tibia length. (n=4-12). Data are means ± SEM; *p<0.05, **p<0.01, ***p<0.001, ****p<0.0001. HYA – Healthy Young Adult, eAMD – early Age-related Muscle Dysfunction, aAMD – advanced Age-related Muscle Dysfunction.

**Supplementary figure 2.**
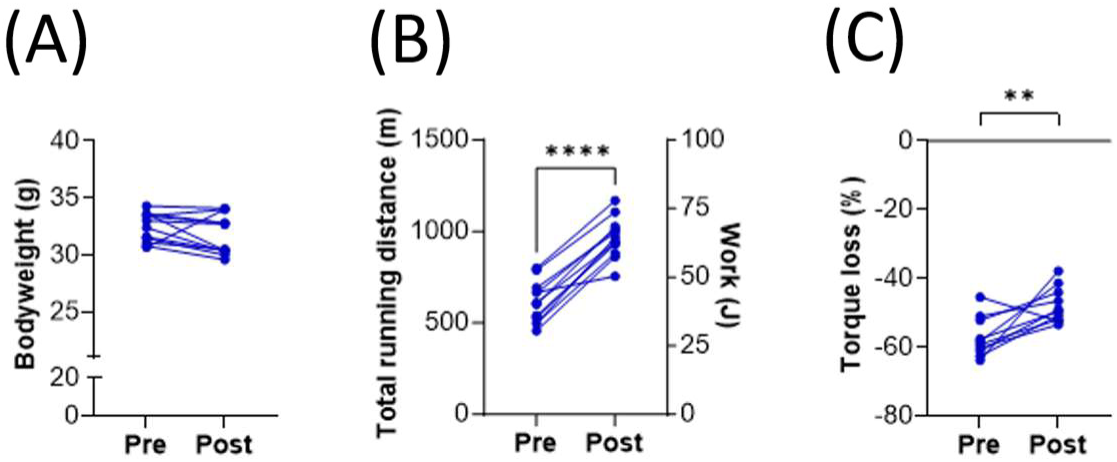
Effects of regular endurance exercise on 21-month-old mice (eAMD+Ex). (A) Bodyweight before and after 6-to-8 week treadmill intervention (n=12). (B) Total running distance during treadmill exhaustion test (n=12). (C) Percentage of initial force lost after 70 repetitive submaximal (50Hz) stimulations. (n=12). Pre – before exercise regular exercise, Post – after regular exercise. Data are individual values; **p<0.01, ****p<0.0001.

**Supplementary figure 3.**
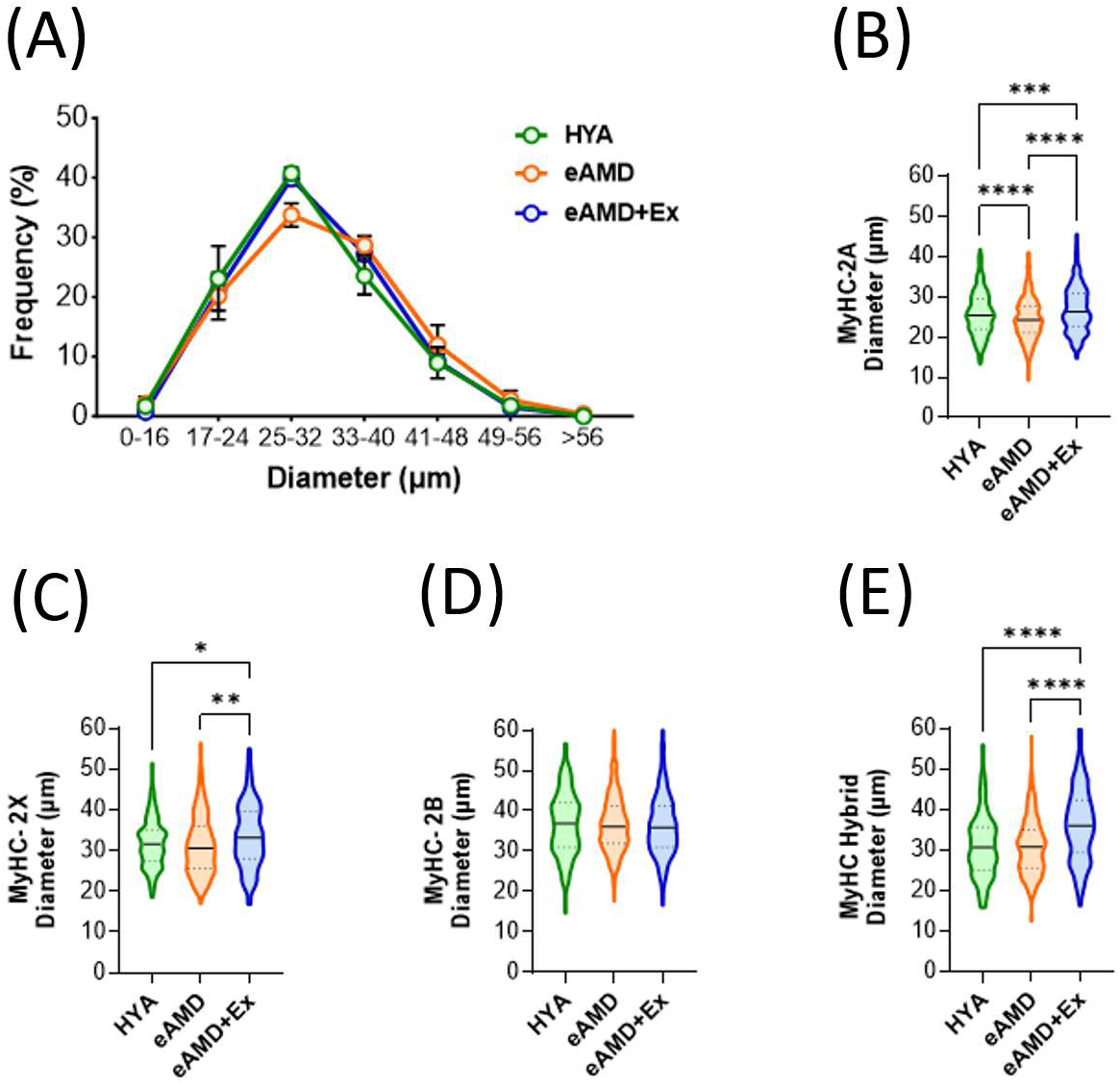
Effects of aging and exercise on plantaris fiber size. (A) Distribution of all fiber diameters across groups. Data are means ± SEM (n=4). (B-E) Diameter of individual fiber types (MyHC-2A, MyHC-2X, MyHC-2B, MyHC-Hybrid) across groups. Violin plots display 1st quartile, median, and 3rd quartile. *p<0.05, **p<0.01, ***p<0.001, ****p<0.0001. HYA – Healthy Young Adult, eAMD – early Age-related Muscle Dysfunction, eAMD+Ex – early Age-related Muscle Dysfunction following 6-8 weeks of regular endurance exercise.

**Supplementary Figure 4.**
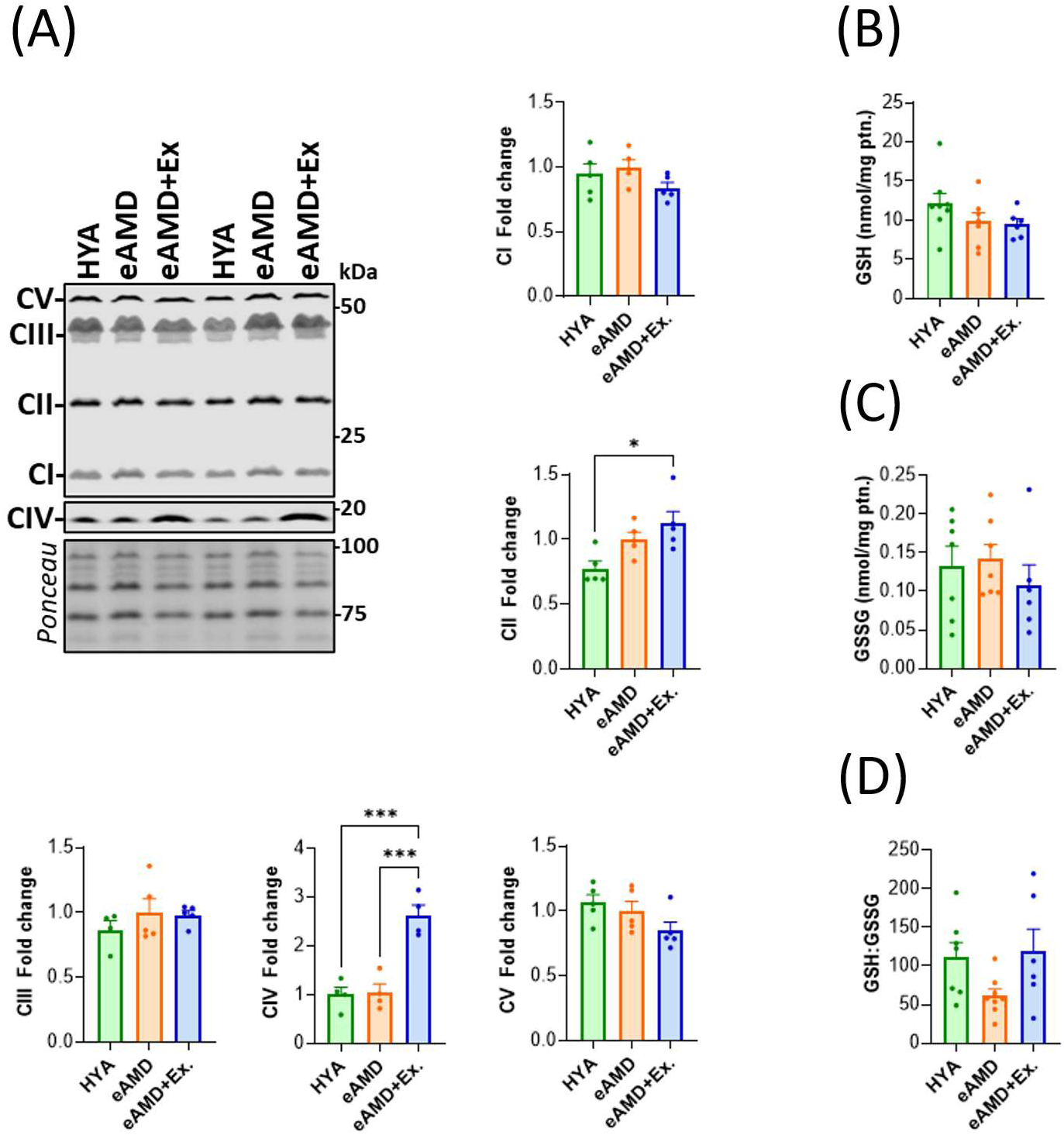
Effects of aging and exercise on skeletal muscle mitochondria. (A) Representative immunoblot of ETC complex units: ATP5A (CV), UQCRC2 (CIII), SDHB (CII), NDUFB8 (CI), and COX IV (CIV) and quantification for each group. Proteins were normalized to Ponceau signal. (N=4). (B) Total concentration of reduced glutathione (GSH) in GA lysates. Values normalized to milligrams of protein per well (n=6-7). (C) Total concentration of oxidized glutathione (GSSG) in GA lysates. Values normalized to milligrams of protein per well (n=6-7). (D) Ratio of GSH concentration to GSSG concentration in each sample (ANOVA p=0.089). Data are means ± SEM; *p<0.05, ***p<0.001. HYA – Healthy Young Adult, eAMD – early Age-related Muscle Dysfunction, eAMD+Ex – early Age-related Muscle Dysfunction following 6-8 weeks of regular endurance exercise.

**Supplementary Figure 5.**
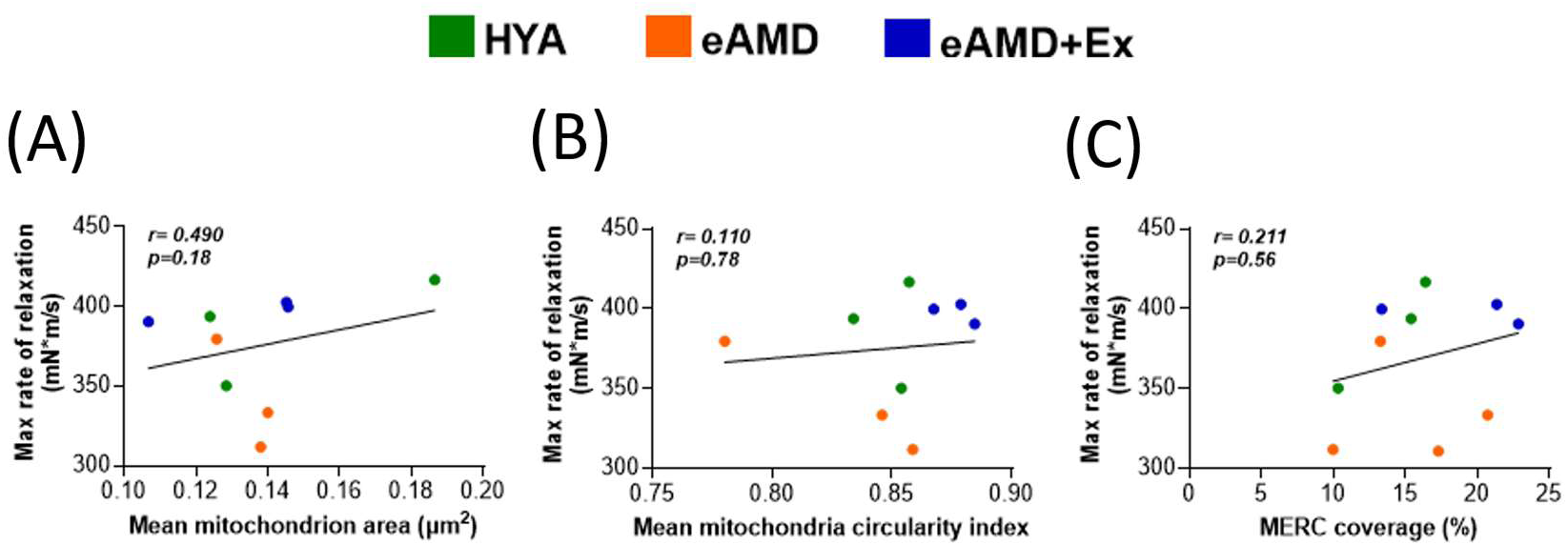
Correlation analyses of significantly-altered ultrastructural parameters and *in vivo* rate of muscle relaxation. Pearson correlation coefficient (*r*) resulting from (A) Average mitochondrial area and relaxation, (B) Average mitochondrial circularity and relaxation, and (C) MERC coverage and relaxation. (n=3/group). HYA – Healthy young adult, eAMD –Early age-related muscle dysfunction, eAMD+Ex – Early age-related muscle dysfunction following 6-8 weeks of regular endurance exercise.

## References

Aagaard, P., Suetta, C., Caserotti, P., Magnusson, S. P., & Kjær, M. (2010). Role of the nervous system in sarcopenia and muscle atrophy with aging: strength training as a countermeasure. Scandinavian Journal of Medicine & Science in Sports, 20(1), 49–64. 10.1111/j.1600-0838.2009.01084.x

Anderson, E. J., Yamazaki, H., & Neufer, P. D. (2007). Induction of endogenous uncoupling protein 3 suppresses mitochondrial oxidant emission during fatty acid-supported respiration. J Biol Chem, 282(43), 31257–31266. doi:10.1074/jbc.M706129200

Andersson, D. C., Betzenhauser, M. J., Reiken, S., Meli, A. C., Umanskaya, A., Xie, W., . . .Marks, A. R. (2011). Ryanodine receptor oxidation causes intracellular calcium leak and muscle weakness in aging. Cell Metab, 14(2), 196–207. doi:10.1016/j.cmet.2011.05.014

Baehr, L. M., West, D. W., Marcotte, G., Marshall, A. G., De Sousa, L. G., Baar, K., & Bodine, S. C. (2016). Age-related deficits in skeletal muscle recovery following disuse are associated with neuromuscular junction instability and ER stress, not impaired protein synthesis. Aging (Albany NY), 8(1), 127–146. doi:10.18632/aging.100879

Bohnert, K. R., McMillan, J. D., & Kumar, A. (2018). Emerging roles of ER stress and unfolded protein response pathways in skeletal muscle health and disease. J Cell Physiol, 233(1), 67–78. doi:10.1002/jcp.25852

Boudina, S., Sena, S., Theobald, H., Sheng, X., Wright, J. J., Hu, X. X., . . .Abel, E. D. (2007). Mitochondrial energetics in the heart in obesity-related diabetes: direct evidence for increased uncoupled respiration and activation of uncoupling proteins. Diabetes, 56(10), 2457–2466. doi:10.2337/db07-0481

Budui, S. L., Rossi, A. P., & Zamboni, M. (2015). The pathogenetic bases of sarcopenia. Clin Cases Miner Bone Metab, 12(1), 22–26. doi:10.11138/ccmbm/2015.12.1.022

Carreras-Sureda, A., Jaña, F., Urra, H., Durand, S., Mortenson, D. E., Sagredo, A., . . .Hetz, C. (2019). Non-canonical function of IRE1α determines mitochondria-associated endoplasmic reticulum composition to control calcium transfer and bioenergetics. Nat Cell Biol, 21(6), 755–767. doi:10.1038/s41556-019-0329-y

Cartee, G. D., Hepple, R. T., Bamman, M. M., & Zierath, J. R. (2016). Exercise Promotes Healthy Aging of Skeletal Muscle. Cell Metab, 23(6), 1034–1047. doi:10.1016/j.cmet.2016.05.007

Chatterji, S., Byles, J., Cutler, D., Seeman, T., & Verdes, E. (2015). Health, functioning, and disability in older adults—present status and future implications. The Lancet, 385(9967), 563–575. 10.1016/S0140-6736(14)61462-8

Cruz-Jentoft, A. J., Landi, F., Schneider, S. M., Zúñiga, C., Arai, H., Boirie, Y., . . .Cederholm, T. (2014). Prevalence of and interventions for sarcopenia in ageing adults: a systematic review. Report of the International Sarcopenia Initiative (EWGSOP and IWGS). Age Ageing, 43(6), 748–759. doi:10.1093/ageing/afu115

Doherty, T. J. (2003). Invited Review: Aging and sarcopenia. Journal of Applied Physiology, 95(4), 1717–1727. doi:10.1152/japplphysiol.00347.2003

Dowling, P., Gargan, S., Swandulla, D., & Ohlendieck, K. (2023). Fiber-Type Shifting in Sarcopenia of Old Age: Proteomic Profiling of the Contractile Apparatus of Skeletal Muscles. International Journal of Molecular Sciences, 24(3), 2415. Retrieved from https://www.mdpi.com/1422-0067/24/3/2415

Duchen, M. R. (2000). Mitochondria and calcium: from cell signalling to cell death. J Physiol, 529 Pt 1(Pt 1), 57–68. doi:10.1111/j.1469-7793.2000.00057.x

Erdjument-Bromage, H., Huang, F. K., & Neubert, T. A. (2018). Sample Preparation for Relative Quantitation of Proteins Using Tandem Mass Tags (TMT) and Mass Spectrometry (MS). Methods Mol Biol, 1741, 135–149. doi:10.1007/978-1-4939-7659-1_11

Fuqua, J. D., Mere, C. P., Kronemberger, A., Blomme, J., Bae, D., Turner, K. D., . . .Lira, V. A. (2019). ULK2 is essential for degradation of ubiquitinated protein aggregates and homeostasis in skeletal muscle. The FASEB Journal, 33(11), 11735–12745. 10.1096/fj.201900766R

Giacomello, M., & Pellegrini, L. (2016). The coming of age of the mitochondria–ER contact: a matter of thickness. Cell Death & Differentiation, 23(9), 1417–1427. doi:10.1038/cdd.2016.52

Hamasaki, M., Furuta, N., Matsuda, A., Nezu, A., Yamamoto, A., Fujita, N., . . .Yoshimori, T. (2013). Autophagosomes form at ER–mitochondria contact sites. Nature, 495(7441), 389–393. doi:10.1038/nature11910

Harris, M. P., Zhang, Q. J., Cochran, C. T., Ponce, J., Alexander, S., Kronemberger, A., . . .Lira, V. A. (2022). Perinatal versus adult loss of ULK1 and ULK2 distinctly influences cardiac autophagy and function. Autophagy, 18(9), 2161–2177. doi:10.1080/15548627.2021.2022289

Hinton, A., Katti, P., Mungai, M., Hall, D., Koval, O., Shao, J., . . .Abel, E. D. (2022). OPA1 Downregulation in Skeletal Muscle Induces MERC formation in an ATF4-Dependent Manner. bioRxiv, 2022.2009.2012.507669. doi:10.1101/2022.09.12.507669

Kauppila, T. E. S., Kauppila, J. H. K., & Larsson, N.-G. (2017). Mammalian Mitochondria and Aging: An Update. Cell Metabolism, 25(1), 57–71. 10.1016/j.cmet.2016.09.017

Kerr, H. L., Krumm, K., Anderson, B., Christiani, A., Strait, L., Li, T., . . .Garcia, J. M. (2024). Mouse sarcopenia model reveals sex- and age-specific differences in phenotypic and molecular characteristics. J Clin Invest, 134(16). doi:10.1172/jci172890

Kubat, G. B., Bouhamida, E., Ulger, O., Turkel, I., Pedriali, G., Ramaccini, D., . . .Pinton, P. (2023). Mitochondrial dysfunction and skeletal muscle atrophy: Causes, mechanisms, and treatment strategies. Mitochondrion, 72, 33–58. doi:10.1016/j.mito.2023.07.003

Lam, J., Katti, P., Biete, M., Mungai, M., AshShareef, S., Neikirk, K., . . .Hinton, A. (2021). A Universal Approach to Analyzing Transmission Electron Microscopy with ImageJ. Cells, 10(9), 2177. Retrieved from https://www.mdpi.com/2073-4409/10/9/2177

Lang, T., Streeper, T., Cawthon, P., Baldwin, K., Taaffe, D. R., & Harris, T. B. (2010). Sarcopenia: etiology, clinical consequences, intervention, and assessment. Osteoporosis International, 21(4), 543–559. doi:10.1007/s00198-009-1059-y

Lark, D. S., Torres, M. J., Lin, C. T., Ryan, T. E., Anderson, E. J., & Neufer, P. D. (2016). Direct real-time quantification of mitochondrial oxidative phosphorylation efficiency in permeabilized skeletal muscle myofibers. Am J Physiol Cell Physiol, 311(2), C239–245. doi:10.1152/ajpcell.00124.2016

Larsson, L., Degens, H., Li, M., Salviati, L., Lee, Y. i., Thompson, W., . . .Sandri, M. (2019). Sarcopenia: Aging-Related Loss of Muscle Mass and Function. Physiological Reviews, 99(1), 427–511. doi:10.1152/physrev.00061.2017

Loeb, G. E., & Ghez, C. (2000). The motor unit and muscle action. Principles of neural science, 380(6573), 674–694.

Lu, X., Gong, Y., Hu, W., Mao, Y., Wang, T., Sun, Z., . . .Lai, D. (2022). Ultrastructural and proteomic profiling of mitochondria-associated endoplasmic reticulum membranes reveal aging signatures in striated muscle. Cell Death & Disease, 13(4), 296. doi:10.1038/s41419-022-04746-4

Lund-Johansen, F., de la Rosa Carrillo, D., Mehta, A., Sikorski, K., Inngjerdingen, M., Kalina, T., . . .Stuchly, J. (2016). MetaMass, a tool for meta-analysis of subcellular proteomics data. Nature Methods, 13(10), 837–840. doi:10.1038/nmeth.3967

Marzetti, E., Calvani, R., Coelho-Júnior, H. J., Landi, F., & Picca, A. (2024). Mitochondrial Quantity and Quality in Age-Related Sarcopenia. Int J Mol Sci, 25(4). doi:10.3390/ijms25042052

Mast, I. H., Allard, N. A. E., Ten Haaf, D., Stoffels, A. A. F., Janssen, L., van Hees, H. W. H., . . .Buffart, L. M. (2024). Muscle contractile properties and perceived fatigue in the general and diseased population. Physiol Rep, 12(23), e70134. doi:10.14814/phy2.70134

Mayfield, D. L., Cronin, N. J., & Lichtwark, G. A. (2023). Understanding altered contractile properties in advanced age: insights from a systematic muscle modelling approach. Biomechanics and Modeling in Mechanobiology, 22(1), 309–337. doi:10.1007/s10237-022-01651-9

Michelucci, A., Liang, C., Protasi, F., & Dirksen, R. T. (2021). Altered Ca2+ Handling and Oxidative Stress Underlie Mitochondrial Damage and Skeletal Muscle Dysfunction in Aging and Disease. Metabolites, 11(7), 424. Retrieved from https://www.mdpi.com/2218-1989/11/7/424

Migliavacca, E., Tay, S. K. H., Patel, H. P., Sonntag, T., Civiletto, G., McFarlane, C., . . .Feige, J. N. (2019). Mitochondrial oxidative capacity and NAD(+) biosynthesis are reduced in human sarcopenia across ethnicities. Nat Commun, 10(1), 5808. doi:10.1038/s41467-019-13694-1

Murphy, A. N., Bredesen, D. E., Cortopassi, G., Wang, E., & Fiskum, G. (1996). Bcl-2 potentiates the maximal calcium uptake capacity of neural cell mitochondria. Proc Natl Acad Sci U S A, 93(18), 9893–9898. doi:10.1073/pnas.93.18.9893

Paez, H. G., Pitzer, C. R., & Alway, S. E. (2023). Age-Related Dysfunction in Proteostasis and Cellular Quality Control in the Development of Sarcopenia. Cells, 12(2). doi:10.3390/cells12020249

Parry, H. A., Roberts, M. D., & Kavazis, A. N. (2020). Human Skeletal Muscle Mitochondrial Adaptations Following Resistance Exercise Training. Int J Sports Med, 41(6), 349–359. doi:10.1055/a-1121-7851

Poston, C. N., Krishnan, S. C., & Bazemore-Walker, C. R. (2013). In-depth proteomic analysis of mammalian mitochondria-associated membranes (MAM). Journal of Proteomics, 79, 219–230. 10.1016/j.jprot.2012.12.018

Powers, S. K., Ji, L. L., Kavazis, A. N., & Jackson, M. J. (2011). Reactive oxygen species: impact on skeletal muscle. Compr Physiol, 1(2), 941–969. doi:10.1002/cphy.c100054

Rappsilber, J., Mann, M., & Ishihama, Y. (2007). Protocol for micro-purification, enrichment, pre-fractionation and storage of peptides for proteomics using StageTips. Nat Protoc, 2(8), 1896–1906. doi:10.1038/nprot.2007.261

Rosencrans, W. M., Rajendran, M., Bezrukov, S. M., & Rostovtseva, T. K. (2021). VDAC regulation of mitochondrial calcium flux: From channel biophysics to disease. Cell Calcium, 94, 102356. 10.1016/j.ceca.2021.102356

Sayer, A. A., Cooper, R., Arai, H., Cawthon, P. M., Ntsama Essomba, M.-J., Fielding, R. A., . . .Cruz-Jentoft, A. J. (2024). Sarcopenia. Nature Reviews Disease Primers, 10(1), 68. doi:10.1038/s41572-024-00550-w

Schefer, V., & Talan, M. I. (1996). Oxygen consumption in adult and aged C57BL/6J mice during acute treadmill exercise of different intensity. Experimental Gerontology, 31(3), 387–392. 10.1016/0531-5565(95)02032-2

Schenk, S., Sagendorf, T. J., Many, G. M., Lira, A. K., de Sousa, L. G. O., Bae, D., . . .Group, T. M. S. (2024). Physiological Adaptations to Progressive Endurance Exercise Training in Adult and Aged Rats: Insights from the Molecular Transducers of Physical Activity Consortium (MoTrPAC). Function, 5(4). doi:10.1093/function/zqae014

Schumann, M., Feuerbacher, J. F., Sünkeler, M., Freitag, N., Rønnestad, B. R., Doma, K., & Lundberg, T. R. (2022). Compatibility of Concurrent Aerobic and Strength Training for Skeletal Muscle Size and Function: An Updated Systematic Review and Meta-Analysis. Sports Med, 52(3), 601–612. doi:10.1007/s40279-021-01587-7

Tarnopolsky, M. A., Rennie, C. D., Robertshaw, H. A., Fedak-Tarnopolsky, S. N., Devries, M. C., & Hamadeh, M. J. (2007). Influence of endurance exercise training and sex on intramyocellular lipid and mitochondrial ultrastructure, substrate use, and mitochondrial enzyme activity. American Journal of Physiology-Regulatory, Integrative and Comparative Physiology, 292(3), R1271–R1278. doi:10.1152/ajpregu.00472.2006

Thoudam, T., Ha, C.-M., Leem, J., Chanda, D., Park, J.-S., Kim, H.-J., . . .Lee, I.-K. (2018). PDK4 Augments ER–Mitochondria Contact to Dampen Skeletal Muscle Insulin Signaling During Obesity. Diabetes, 68(3), 571–586. doi:10.2337/db18-0363

Tubbs, E., Chanon, S., Robert, M., Bendridi, N., Bidaux, G., Chauvin, M.-A., . . .Rieusset, J. (2018). Disruption of Mitochondria-Associated Endoplasmic Reticulum Membrane (MAM) Integrity Contributes to Muscle Insulin Resistance in Mice and Humans. Diabetes, 67(4), 636–650. doi:10.2337/db17-0316

Turner, K. D., Kronemberger, A., Bae, D., Bock, J. M., Hughes, W. E., Ueda, K., . . .Lira, V. A. (2022). Effects of Combined Inorganic Nitrate and Nitrite Supplementation on Cardiorespiratory Fitness and Skeletal Muscle Oxidative Capacity in Type 2 Diabetes: A Pilot Randomized Controlled Trial. Nutrients, 14(21), 4479. Retrieved from https://www.mdpi.com/2072-6643/14/21/4479

von Haehling, S., Morley, J. E., & Anker, S. D. (2010). An overview of sarcopenia: facts and numbers on prevalence and clinical impact. J Cachexia Sarcopenia Muscle, 1(2), 129–133. doi:10.1007/s13539-010-0014-2

Wang, H., Huang, W. Y., & Zhao, Y. (2022). Efficacy of Exercise on Muscle Function and Physical Performance in Older Adults with Sarcopenia: An Updated Systematic Review and Meta-Analysis. Int J Environ Res Public Health, 19(13). doi:10.3390/ijerph19138212

Wieckowski, M. R., Giorgi, C., Lebiedzinska, M., Duszynski, J., & Pinton, P. (2009). Isolation of mitochondria-associated membranes and mitochondria from animal tissues and cells. Nature Protocols, 4(11), 1582–1590. doi:10.1038/nprot.2009.151

Wu, T., Hu, E., Xu, S., Chen, M., Guo, P., Dai, Z., . . .Yu, G. (2021). clusterProfiler 4.0: A universal enrichment tool for interpreting omics data. The Innovation, 2(3), 100141. 10.1016/j.xinn.2021.100141

Zecha, J., Satpathy, S., Kanashova, T., Avanessian, S. C., Kane, M. H., Clauser, K. R., . . .Kuster, B. (2019). TMT Labeling for the Masses: A Robust and Cost-efficient, In-solution Labeling Approach *^[S]^. Molecular & Cellular Proteomics, 18(7), 1468–1478. doi:10.1074/mcp.TIR119.001385

Zhou, M., Kong, B., Zhang, X., Xiao, K., Lu, J., Li, W., . . .Xu, T. (2023). A proximity labeling strategy enables proteomic analysis of inter-organelle membrane contacts. iScience, 26(7), 107159. 10.1016/j.isci.2023.107159

Zollinger, D. R., Baalman, K. L., & Rasband, M. N. (2015). The Ins and Outs of Polarized Axonal Domains. Annual Review of Cell and Developmental Biology, 31(Volume 31, 2015), 647–667. 10.1146/annurev-cellbio-100913-013107

