## Supplementary document 8 for "Altered relaxation and Mitochondria-Endoplasmic Reticulum Contacts Precede Major (Mal)adaptations in Aging Skeletal Muscle and are Prevented by Exercise"

**SUPPLEMENTARY METHODS**

**Mitochondrial O_2_ consumption, ATP production, and H_2_O_2_ emission in permeabilized fibers**

ATP production rates were obtained using enzyme-coupled reaction methods with NADPH production as surrogate readout. NADPH fluorescence (340/460 excitation/emission) was detected via a fiber optic cable (Molex, Lisle, IL) to measure ATP production simultaneously with O_2_ consumption. Immediately prior to experiments, ice-cold permeabilized fibers were transferred from their wash in Buffer Z (105 mM K-MES, 30 mM KCl, 10 mM K_2_HPO_4_, 5 mM MgCl_2_ · 6H_2_O, 0.5 mg/mL fatty acid-free BSA [Sigma-Aldrich, A8806], pH 7.4), weighed into ~3 mg bundles and placed in 2.5 mL Respiration Buffer (Buffer Z + 1 mM EGTA + 20 µM blebbistatin [Sigma-Aldrich, B0560]) supplemented with 1 U/ml of Hexokinase, 2.5 U/ml G-6-PDH, 2.5 mM d-glucose, 200 µM Ap5A and 2.5 mM NADP+. Respiration was supported with pyruvate (5 mM), malate (0.5 mM), ADP (125 µM), glutamate (5 mM), and succinate (5 mM). Fatty acid-mediated respiration was supported with 10 mM palmitoyl-carnitine and the same concentrations of malate, ADP, and succinate. New substrates were injected into the chambers following a plateau of oxygen consumption rate from baseline and/or previous addition(s).

Amplex Red reagent reacts with H_2_O_2_ in the presence of horseradish peroxidase (HRP) to synthesize the fluorescent compound resorufin (excitation/emission = 563 nm/587 nm) (Anderson et al, 2007). Cuvettes were filled with 1mL respiration buffer, 10 µM Amplex Red, 1 U/mL HRP, 5 U/mL superoxide dismutase, 2 mM malate, and 5 mM pyruvate. Fibers were weighed immediately upon removal from the respiration buffer and ~ 1.5 mg were placed into individual cuvettes and loaded in a spectrofluorometer (Photon Technology Instruments, Birmingham, NJ). Experiments were run at 30 °C and began with two to four minutes of background fluorescence (ΔF/min) readings in the presence of pyruvate/malate, after which 5 mM succinate was added.

**Isolation of mitochondrial-associated ER membranes (MAMs)**

After a brief wash in starting buffer (225 mM mannitol, 75 mM sucrose, and 30 mM Tris–HCl, pH 7.4), GAs from both legs were homogenized together using a Wheaton Overhead Stirrer (DWK Life Sciences, Millville, NJ) in Potter-Elvehjem glass tubes containing 2 mL of buffer 1 (225 mM mannitol, 75 mM sucrose, 0.5 % BSA, 0.5 mM EGTA and 30 mM Tris–HCl, pH 7.4). Homogenates were centrifuged at 800 x *g* for 5 min at 4 °C to remove unbroken cells and nuclei. The supernatant was collected, and the previous centrifugation was repeated. Once again, this supernatant was collected, and the sample was then centrifuged at 9000 x *g* for 10 minutes at 4 °C. Upon discarding the new supernatant, the mitochondrial pellet was resuspended by hand using small Teflon pestles in a Potter-Elvehjem glass tube containing 1 mL of starting buffer. These two steps (i.e., centrifugation and resuspension) were repeated twice more to obtain a pellet containing the ‘purified’ crude mitochondrial fraction. Due to small mitochondrial pellets, we used 2 mL Eppendorf tubes and carried out the early centrifugation steps within a ThermoFisher Fresco 21^TM^ microcentrifuge (ThermoFisher, Waltham, MA). The pellet was resuspended in 2 mL of mitochondrial resuspension buffer (MRB- 250 mM mannitol, 5 mM HEPES (pH 7.4) and 0.5 mM EGTA) then layered on top of a Percoll density gradient (225 mM mannitol, 25 mM HEPES (pH 7.4), 1 mM EGTA and 30 % Percoll (vol/vol)) and subjected to 95000 x *g* for 30 min at 4 °C. This step then effectively separated ‘pure mitochondria’ and ‘mitochondria-associated membranes’ from each other in two visible bands (Lu et al, 2022). The latter was collected into a 1.5 mL tube and centrifuged at 6300 x *g* for 10 min at 4 °C. To obtain a final MAM pellet, the supernatants were centrifuged at 100000 x *g* for 1 hour at 4 °C. Protease (Roche cOmplete^TM^, mini, EDTA-free) and phosphatase (Roche PhosSTOP^TM^) inhibitors cocktail tablets were added to the final samples.

**Proteomics samples preparation**

An estimated 1.25 ug of each sample was loaded on NuPage 4-12 % Bis-Tris precast 15-well gel (Invitrogen, USA) and separated at 200 V for 50 minutes. 10 μL solution of mass ladder markers (Sharp pre-stained migration standards, Invitrogen) was loaded onto a separate gel lane to serve as a guide to molecular weight. The gel was stained using a Pierce mass spec compatible silver stain kit (Thermo Scientific, USA) following the manufacturer’s directions.

Proteins were initially quantified by a micro-BCA assay (ThermoFisher 23235). Six micrograms of each sample were reduced and alkylated in 100 µl lysis buffer (6 M Guanidinium hydrochloride (Gdn), 10 mM DTT) at 56 °C for 1 hour, then 50 mM chloroacetamide (CAA), 100 mM Tris–HCl (pH 8.5) in the dark at room temperature for 30 min. After cooling to room temperature, proteins were digested with LysC (Wako) at the ratio of enzyme-to-protein of 1:30 (w/w) at 37 °C for 5 hours. Samples were diluted with 25mM Tris-HCl pH 8.5 to a final concentration 1M in GdnHCl, then incubated with trypsin at a 1:20 ratio overnight (14 hours). The next morning, additional trypsin was added to each sample to establish a 1:50 protease to protein ratio (w/w) and digestion continued for 3 hours at 37 °C. Digested samples were acidified to pH 2-3 with 50 % TFA (trifluoroacetic acid) and centrifuged at 20,000 x g for 15 min to pellet insoluble material (Rappsilber et al, 2007). The supernatant peptides were desalted with C18 stage tips Peptides were eluted in 200 µl 70 % ACN and 0.1 % FA. Supernatants were checked by MALDI-TOF, and the remainder was concentrated by lyophilization and then reconstituted in 20 µL of mobile phase A (0.1 % formic acid with 3 % acetonitrile) and stored at 20 °C for further analyses.

**MALDI-TOF**

Each sample (2 μL) was mixed with the same volume of saturated HCCA matrix (Bruker, Billerica, MA, USA) in 50 % acetonitrile and applied to 384-well Anchor Chip plates (Bruker) then air dried. The plate was mechanically transported into the source vacuum of a Bruker MALDI-TOF/TOF FlexeXtreme and approximately 2k from a 355 nm laser were acquired in reflectron mode shots per spectrum covering a mass range from 700 Da to 5k Da. Supplementary Figure 9.1 (shown below) illustrates a distinctive pattern for polyvinyl pyrrolidone (PVP) and polyethylene glycol (PEG). PVP and PEG components are identified by a recurring pattern of peaks at 111.1 Da and 44.06 Da, respectively. MAMs were isolated on a density gradient of Percoll (Cytiva, Wilmington, DE, USA), which consists of PVP molecularly bonded to silica particles,15-30 μm in diameter. The manufacturer’s usage guide states that free PVP makes up to 2% of the total reagent. Free PVP would cause peptide suppression and extreme carryover between Liquid Chromatography injections and may remodel capillary packing. Thus, PVP is likely to interfere with differential expression profiling among samples. To combat this, we used a modification of a gel-assisted method reported by which MAM isolates are entrapped in a polyacrylamide gel matrix and wshed thoroughly with no electrophoresis taking place before digestion is performed within the gel network (Poston et al, 2013). Based on BCA assays, all samples were concentrated to 0.338 μg/μL. Handling each sample individually, a single drop polyacrylamide gel was created by adding 37 μL of 40 % Acrylamide, 5 μL of 10 % Ammonium Persulfate and 2 μL of TEMED to 12 μL of sample in a 1.5 mL Axygen snap cap tube. These were stored at 4 °C for 2 h, and then cut into small 1.5 mm pieces. The gel pieces were washed 3 times with 400 μL of fresh 50 % acetonitrile and 50 % 25 mM NH_4_HCO_3_ (pH 8.0). The gels were dried in 100 % ACN. The supernatant was replaced with 10 mM DTT for reduction at 56 °C for 1 h and the samples were made 50 mM in CAA and left in the dark to alkylate for 30 minutes. Again, the gel pieces were washed 3 times using 400 μ L of fresh 50% ACN in 25 mM NH_4_HCO_3_. Next, trypsin went in 25 mM NH_4_HCO_3_ (pH=8) to digest overnight at 37 °C. Three extracts of the gel pieces with 60 % ACN in 0.1 % formic acid were combined for each sample. These extracts were desalted with stage tips, the elute frozen, lyophilized, and then stored at -80 °C.

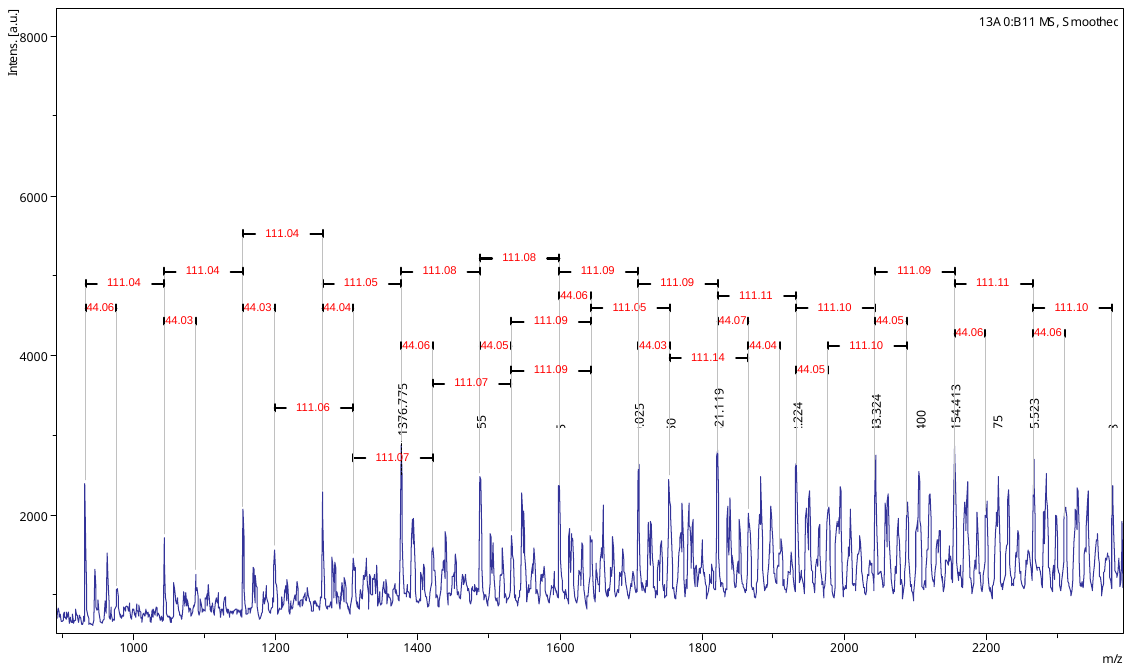

**Supplementary Figure 9.1**. MS1 MALDI spectra of in-solution digest of equal sample mixture

**References**

Anderson, E. J., Yamazaki, H., & Neufer, P. D. (2007). Induction of endogenous uncoupling protein 3 suppresses mitochondrial oxidant emission during fatty acid-supported respiration. *J Biol Chem, 282*(43), 31257-31266. doi:10.1074/jbc.M706129200

Lu, X., Gong, Y., Hu, W., Mao, Y., Wang, T., Sun, Z., . . .Lai, D. (2022). Ultrastructural and proteomic profiling of mitochondria-associated endoplasmic reticulum membranes reveal aging signatures in striated muscle. *Cell Death & Disease, 13*(4), 296. doi:10.1038/s41419-022-04746-4

Poston, C. N., Krishnan, S. C., & Bazemore-Walker, C. R. (2013). In-depth proteomic analysis of mammalian mitochondria-associated membranes (MAM). *Journal of Proteomics, 79*, 219-230. doi:<https://doi.org/10.1016/j.jprot.2012.12.018>

Rappsilber, J., Mann, M., & Ishihama, Y. (2007). Protocol for micro-purification, enrichment, pre-fractionation and storage of peptides for proteomics using StageTips. *Nat Protoc, 2*(8), 1896-1906. doi:10.1038/nprot.2007.261
